# The adaptive architecture of tRNA dependencies across physiological tumor environments

**DOI:** 10.64898/2026.10.08.757597

**Authors:** Siyu Chen, Trey Charbonneau, Kian Yousefi, Bahar Zirak, Sean Lee, Tanvi Joshi, April Pawluk, Vijay Ramani, Hani Goodarzi

**Author notes:** Corresponding authors: H.G., V.R.

## Abstract

The genetic code is decoded by a highly redundant transfer RNA (tRNA) repertoire, yet whether this apparent redundancy serves a functional role beyond ensuring robust translation remains unclear. Here, we establish an atlas of tRNA dependencies through large-scale CRISPRi perturbation mapping across matched *in vivo* tumor growth and *in vitro* cell culture for 16 cancer cell lines spanning seven tissue types. During *in vivo* tumor growth, tRNA dependencies exhibited tissue organization at isodecoder resolution and were associated with codon demand at the isoacceptor level. Strikingly, these relationships were reorganized when cells were cultured *in vitro*, establishing environmental sensitivity of tRNA dependency–in contrast to protein components of the decoding machinery such as aminoacyl-tRNA synthetases (aaRSs), whose dependencies were largely preserved across environments. Despite pronounced context specificity, environmental alterations in tRNA dependency were dominated by a shared response across cancer cell lines, organized across isoacceptor and isodecoder levels. By disentangling stable cellular identity from environment-associated dependency states, we resolved a shared axis of isoacceptor dependency remodeling and identified specific tRNA isoacceptors associated with promoting or restraining adaptation to the tumor environment. We further connect this functional architecture to differential translation of codon-usage-biased programs associated with proliferation and tumor-microenvironmental stress. Together, our study reveals that tRNA redundancy does not imply functional equivalence, but instead forms a structured and environmentally plastic control layer linking coding information to translational output.

## 1. Introduction

The functional output of the genome depends not only on protein-coding genes but also on a vast repertoire of noncoding DNA and RNA elements that regulate when, where, and how genetic information is expressed (King and Wilson, 1975; Wittkopp and Kalay, 2011). Disease-associated genetic variation is highly enriched within noncoding regulatory sequences, and recent studies have increasingly implicated both DNA regulatory elements (Maurano et al., 2012; Emani et al., 2024; Avsec et al., 2026; Kellman et al., 2025) and noncoding RNAs (Rius et al., 2026; De Jonghe et al., 2026; Greene et al., 2024; Lee et al., 2025; Moir et al., 2024; Merheb et al., 2021; Zhu et al., 2026) in human disease. Many classes of noncoding elements are organized as multi-copy or functionally overlapping repertoires (Goodenbour and Pan, 2006; Alvarez-Saavedra and Horvitz, 2010; Osterwalder et al., 2018), act through sequence- and context-dependent interactions, and can acquire markedly different functional requirements across cellular states and environments (Namba et al., 2026; Li and Zhang, 2018). Defining the principles that govern their context-dependent roles therefore represents a major challenge in understanding noncoding genome function.

Transfer RNAs (tRNAs) provide a particularly striking example of this problem. The human genome encodes hundreds of tRNA genes that collectively form a highly redundant decoding system: multiple tRNA genes can generate identical or closely related mature molecules, multiple isodecoders can decode the same codon, and multiple isoacceptors can carry the same amino acid while decoding different synonymous codons (Goodenbour and Pan, 2006). This redundancy has traditionally been viewed as supporting robust and efficient translation through the matching of tRNA supply to codon demand (Ikemura, 1981, 1985; dos Reis et al., 2004; Akashi, 1994). However, accumulating evidence indicates that the tRNA repertoire is actively regulated rather than functionally interchangeable (Pinzaru and Tavazoie, 2023). tRNA transcription by RNA polymerase III responds to transcriptional and nutrient-sensitive regulatory pathways (Willis and Moir, 2018; Lee et al., 2012), while the mature tRNA pool is further shaped by changes in abundance, modification, aminoacylation, and turnover. tRNAs can also give rise to tRNA-derived fragments with regulatory functions beyond translation (Goodarzi et al., 2015), although the extent and diversity of these fragments across cell types remain debated (Gustafsson et al., 2025). The apparent redundancy of the tRNA repertoire therefore appears to mask substantial molecular and functional specialization, yet which tRNAs are functionally required across distinct cellular and physiological states remains poorly understood.

Understanding the functional organization of the tRNA repertoire is central to understanding how genetic information is translated into protein output. It remains challenging to predict protein production from genome sequence or transcript abundance, reflecting the dynamic and incompletely understood nature of post-transcriptional and translational control across cellular states and physiological environments (Buccitelli and Selbach, 2020; Chen et al., 2024; Yue et al., 2026; Tretyachenko et al., 2026; Tsour et al., 2026). As a central component of the machinery that interprets genetic codons, tRNAs provide a direct interface between coding information and translation. Changes in tRNA composition can reshape codon-specific translation and have been linked to cell-state transitions, tumor initiation, metastatic progression, and other disease-associated phenotypes (Goodarzi et al., 2016; Pinzaru and Tavazoie, 2023; Passarelli et al., 2022; Stanley et al., 2026; Pataskar et al., 2022). However, whether the functional requirements of the decoding machinery itself change across physiological environments remains largely unknown. Resolving this question requires moving beyond measurements of tRNA abundance or molecular state to directly determine which components of the decoding machinery are functionally required in different contexts, an approach enabled by genetic dependency mapping (Gilbert et al., 2014; Tsherniak et al., 2017). This is particularly relevant in cancer, where tumor cells encounter nutrient limitation, hypoxia, extracellular signals, and other microenvironmental pressures that reshape metabolic and translational demands (Hanahan, 2022, 2026; Pavlova and Thompson, 2016; Biffo et al., 2024; Fabbri et al., 2021). Mapping how tRNA requirements vary with cellular identity and environmental context at scale may therefore help reveal how changes in the decoding machinery contribute to context-specific translational and proteomic states, and ultimately to the establishment of cellular state.

Here, we establish a systematic dependency map of the human tRNA repertoire through parallel large-scale CRISPR interference (CRISPRi) screens across 16 cancer cell lines spanning seven tissue types under matched *in vivo* tumor growth and *in vitro* cell culture conditions. We map functional dependencies from individual isodecoders to isoacceptor and amino-acid levels and determine how their organization changes across cellular and environmental contexts. Our results reveal that redundancy does not imply functional equivalence among tRNA species. Rather, tRNA requirements form a hierarchical architecture that is structured and environmentally plastic: during physiological tumor growth, dependencies exhibit tissue-specific organization at the isodecoder level and are associated with codon demand at the isoacceptor level, while changes between environments are dominated by a shared response that is differentially manifested across individual tRNAs and cellular contexts. This environmental sensitivity distinguishes tRNAs from the comparatively stable dependencies of aminoacyl-tRNA synthetases (aaRSs). Disentangling stable cellular identity from environment-associated dependency states further identifies individual isoacceptors that are positively or negatively associated with adaptation to the physiological tumor environment, which we term plasticity drivers and plasticity restrainers. Integration with human tumor multi-omics and independent ribosome-profiling data further connects this functional architecture to differential translation of codon-usage-biased proliferation and stress-associated programs. Together, these findings reveal that the apparent redundancy of the human tRNA repertoire encodes a context-dependent functional organization, forming a dynamic decoding layer through which proteomic output can adapt to environmental conditions.

## 2. Results

### 2.1 A pan-cancer *in vivo* CRISPRi atlas reveals global and tissue-specific tRNA dependencies during tumor growth

To define how the human tRNA repertoire supports tumor growth across diverse physiological contexts, we constructed a pan-cancer *in vivo* CRISPRi atlas by systematically perturbing the human tRNA repertoire across 16 cancer cell lines spanning seven tissue types (**Figure 1A**). This atlas provides a unified view of tRNA-mediated dependencies during tumor growth under physiological selective pressures. Briefly, each human cancer cell line was engineered to stably express ZIM3 KRAB–dCas9 (Alerasool et al., 2020) and transduced with a pooled CRISPRi library targeting tRNA genes, together with non-targeting controls and reference guides targeting aminoacyl-tRNA synthetases (aaRSs) and tRNA-modifying enzymes (see Methods). Following lentiviral transduction (*T*_0_), cells were implanted subcutaneously into NOD scid gamma (NSG) mice, and the resulting xenograft tumors were harvested approximately two weeks later (*T*_*t*_). Perturbation effects were quantified by changes in guide RNA abundance between *T*_0_ and *T*_*t*_ (**Figures 1B** and **S1A**), hereafter referred to as “perturbation scores”.

**Figure 1.**
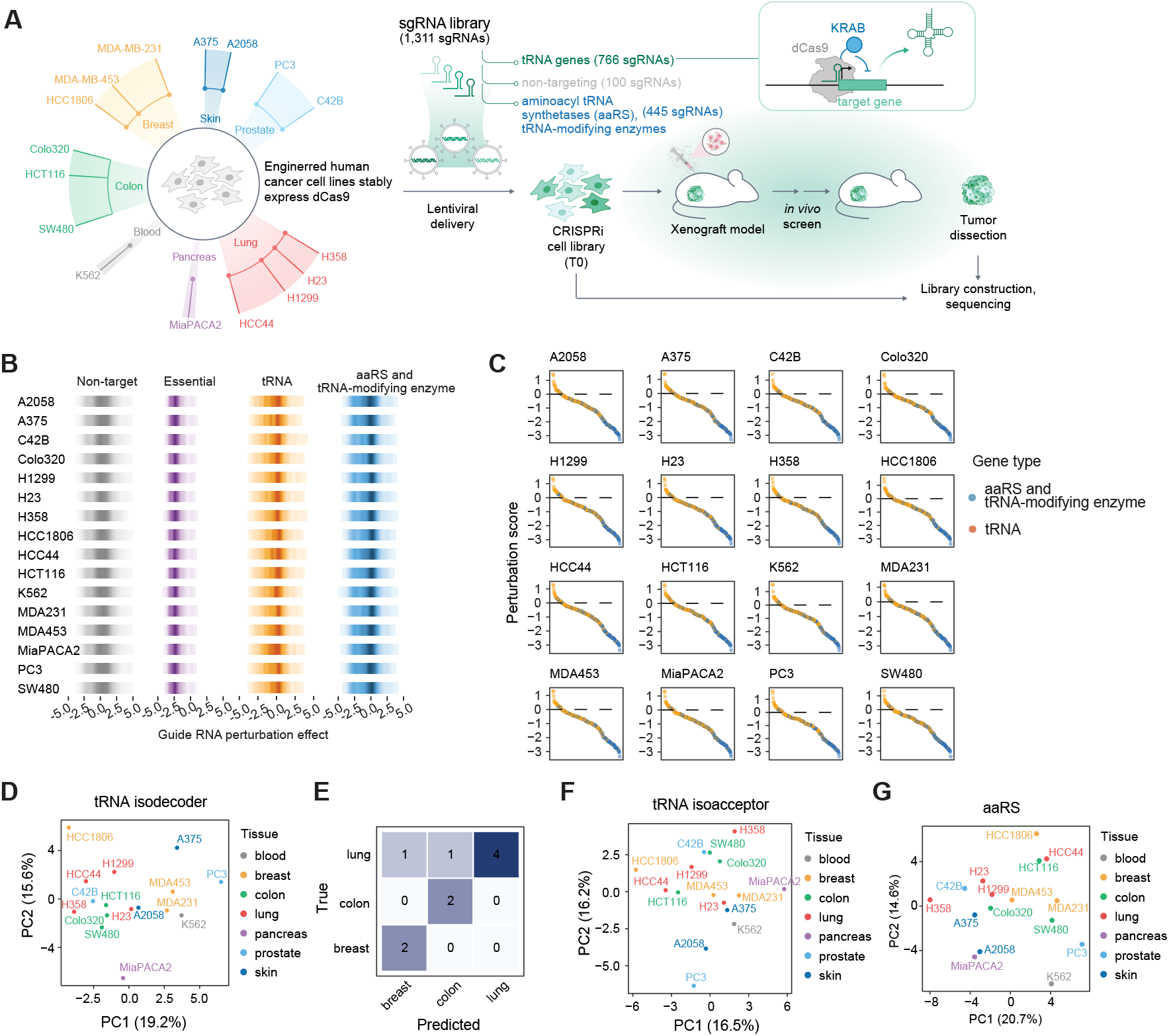
A pan-cancer CRISPRi map of tRNA dependencies during *in vivo* tumor growth. **(A)** Overview of the *in vivo* tRNA CRISPRi atlas encompassing 16 human cancer cell lines across seven tissue types. The single-guide RNA (sgRNA) library comprises 1,311 sgRNAs, including 766 targeting all cytoplasmic tRNA decoding groups (elongator tR-NAs, initiator Met (iMet), and selenocysteine (Sec)), 100 non-targeting controls, and 445 targeting 86 reference genes encoding aminoacyl-tRNA synthetases (aaRSs) and tRNA-modifying enzymes. **(B)** Distribution of guide RNA perturbation effects across individual cell lines. Essential guides refer to guide RNAs targeting commonly essential genes, as defined by DepMap, included in our study. **(C)** Waterfall plots showing perturbation effect distributions of tRNA molecules (orange) and aaRSs and tRNA-modifying enzymes (blue) across cancer cell lines. **(D)** Principal component analysis (PCA) of tRNA isodecoder perturbation effects across cancer cell lines, colored by tissue type. **(E)** Confusion matrix of leave-one-out (LOO) supervised classification showing predicted versus true tissue types based on tRNA isodecoder perturbation landscapes. **(F–G)** PCA plots similar to **(D)** but at the tRNA isoacceptor level **(F)** and for aaRS **(G)**.

To benchmark the performance of our screening approach, we first compared perturbation scores for aaRSs and tRNA-modifying enzymes with corresponding dependency profiles from DepMap (DepMap, Broad, 2024; Arafeh et al., 2025). Despite differences between CRISPR knockout (DepMap) and CRISPRi (this study), perturbation scores were highly concordant across cell lines (Pearson *R* = 0.6; **Figures 1C** and **S1B**), supporting effective CRISPRi perturbation across our independently engineered ZIM3 KRAB–dCas9 cell lines. We separately evaluated the molecular efficiency of tRNA-directed CRISPRi using tRNA-specific RT-qPCR following individual perturbations in MDA-MB-231 triple-negative breast cancer cells, which confirmed substantial depletion of the targeted tRNAs (**Figure S1C**). Together, these complementary validations support effective CRISPRi perturbation across the cell-line panel and efficient molecular depletion of targeted tRNAs, establishing our approach as a robust platform for mapping tRNA and tRNA-related gene dependencies and quantifying their functional consequences during tumor growth.

Having technically validated key aspects of our screening approach, we next asked how perturbing aaRSs, tRNA-modifying enzymes, and individual tRNAs each influenced tumor fitness. As expected, aaRS and tRNA-modifying enzyme perturbations were broadly deleterious, resulting in strongly left-skewed fitness distributions (**Figure 1C**). In contrast, silencing individual tRNA loci produced fitness effects centered near neutrality (**Figure 1C**), indicating that few individual tRNA loci are independently essential for tumor growth. This buffering is consistent with observations in yeast, where deletion of individual tRNA genes is frequently tolerated (Bloom-Ackermann et al., 2014). These results suggest that buffering of individual tRNA loci is evolutionarily conserved and highlight the substantial compensatory capacity of the cellular tRNA pool (Hughes et al., 2023).

Although individual tRNA perturbations were globally buffered, tissue-specific tRNA expression has previously been observed across human cancer cells (Dittmar et al., 2006), raising the possibility that dependency patterns may likewise encode tissue-specific organization. Principal component analysis revealed clear tissue-specific organization at the isodecoder level (*i*.*e*., tRNAs with identical anticodons but differing body sequences; **Figures 1D** and **S2A–B**). We therefore asked whether isodecoder dependency alone was sufficient to classify

**Figure S3.**
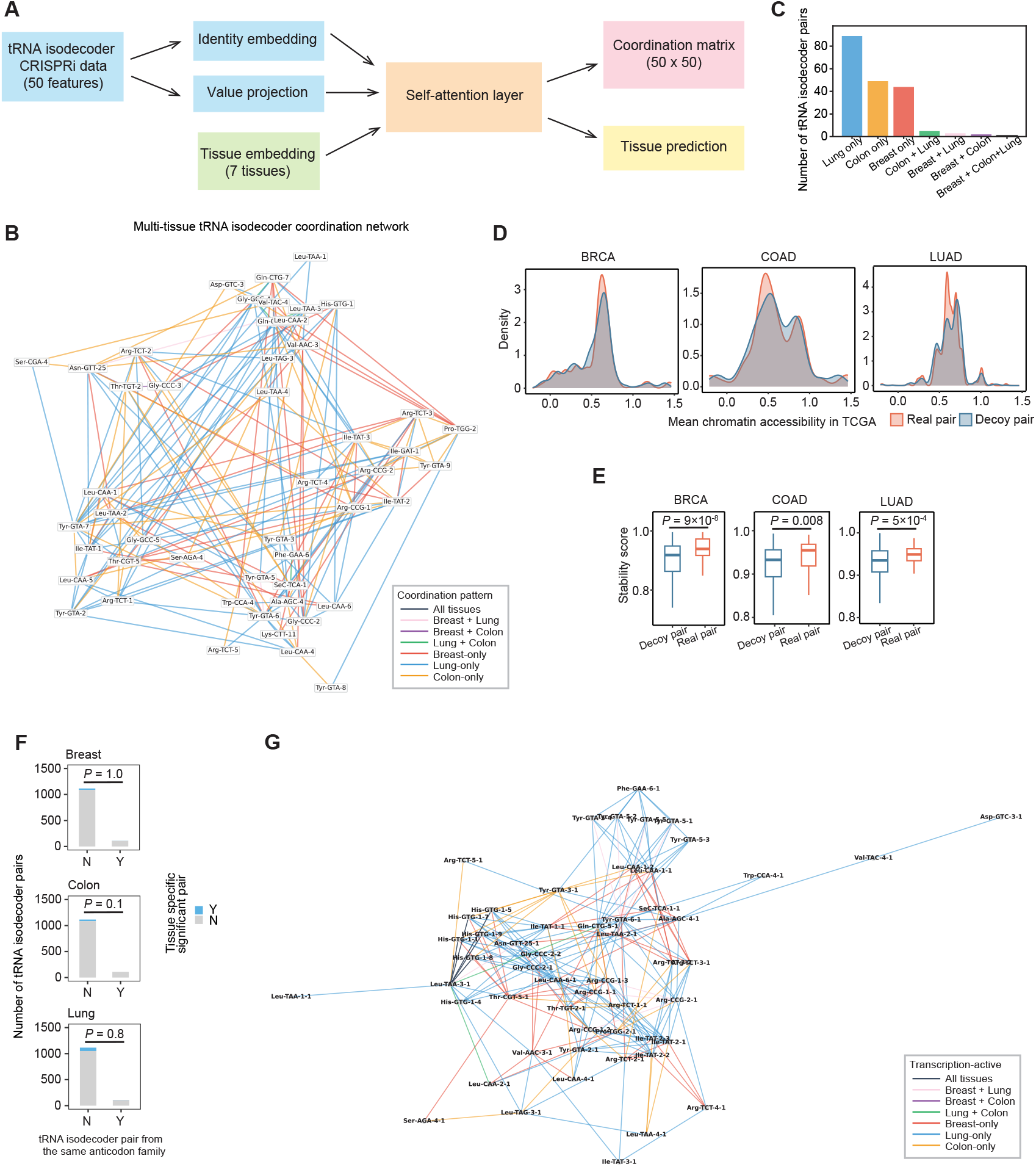
Distinct structures underlie tissue-dependent tRNA isodecoder essentiality. **(A)** Schematic illustration of the multi-tissue self-attention model with contrastive learning used to identify shared and tissue-specific associative structures among tRNA isodecoders. **(B)** Network representation of significant tRNA isodecoder coordination pairs, highlighting shared and tissue-specific modules. Significant pairs were identified by permutation testing (*P <* 0.05) and stability *>* 0.7. **(C)** Bar plot summarizing the numbers of tissue-specific and shared significant tRNA isodecoder pairs shown in **B. (D)** Density plot showing the distribution of chromatin accessibility for real coordination pairs (i, j) and matched chromatin accessibility–controlled decoy partners (j’) derived from TCGA data for the corresponding cancer types. **(E)** Box plot showing the stability of significant real pairs compared with chromatin-matched decoy pairs in the multi-tissue self-attention model. *P*-values were calculated using paired Mann–Whitney *U* tests. **(F)** Bar plot showing the proportion of significant tRNA isodecoder pairs arising from the same amino acid family (Y, blue) or different families (N, gray). *P*-values were calculated using Fisher’s exact test. **(G)** Network representation of active tRNA loci inferred from H3K4me3 ENCODE data, corresponding to the significant tRNA isodecoder coordination pairs in **(B)**.

To rule out the possibility that these tissue-specific isodecoder co-dependencies were driven by differences in CRISPRi efficiency associated with tissue-specific chromatin accessibility, we compared observed isodecoder pairs with chromatin-matched decoy partners using TCGA pan-cancer ATAC-seq data (Corces et al., 2018). For each tissue-specific isodecoder pair (*i, j*), alternative partners *j*^′^ were selected to match *j* in tissue-specific chromatin accessibility (**Figure S3D**). Across breast, colon, and lung, observed isodecoder pairs consistently exhibited stronger dependency coordination than their accessibility-matched decoys (**Figure S3E**; Mann– Whitney *U* tests, *P* = 9 ×10^−8^, 0.008, and 5 ×10^−4^ for breast, colon, and lung, respectively), indicating that the observed coordination cannot be explained by differences in baseline chromatin accessibility alone.

**Figure S1.**
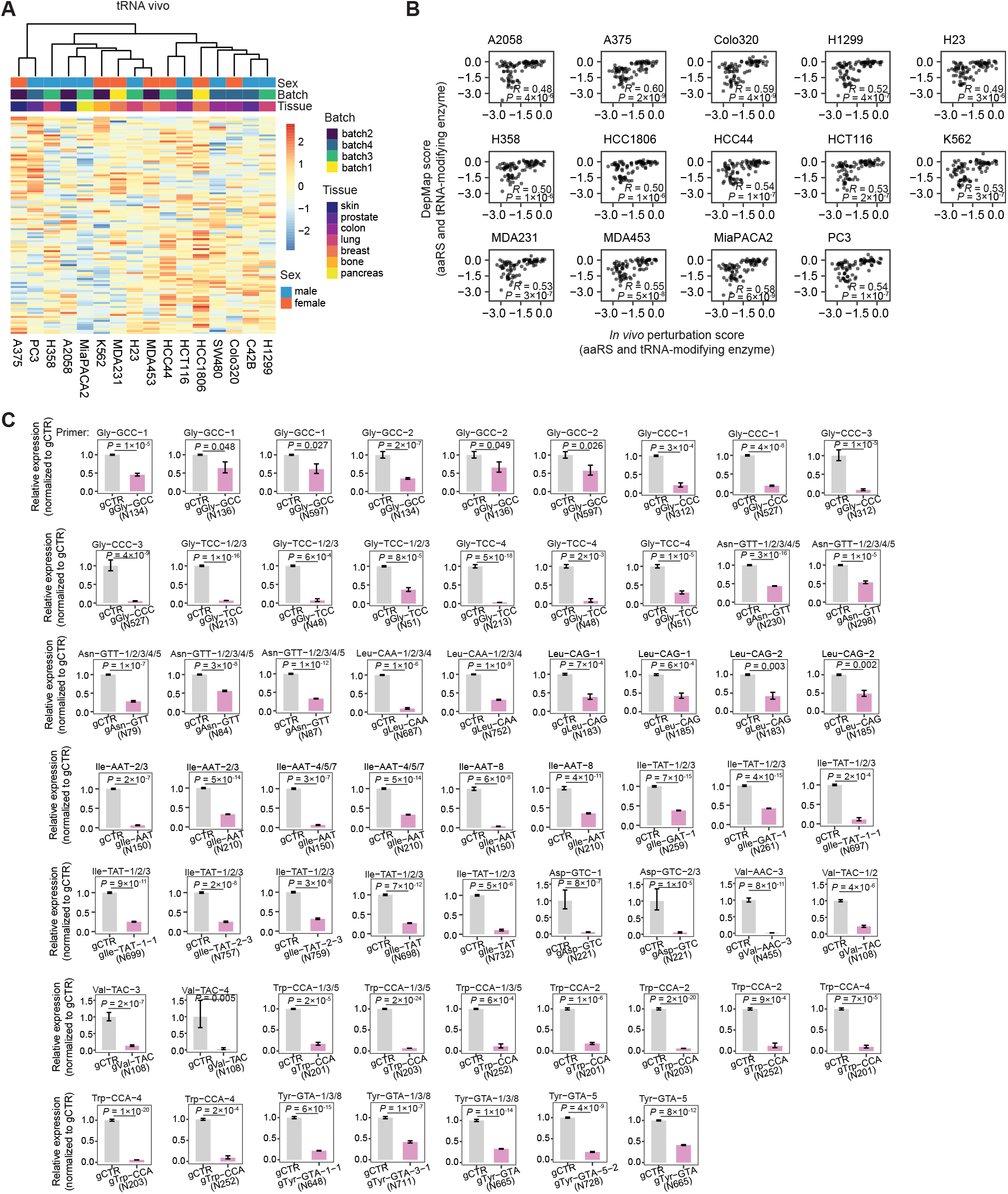
Profiles of *in vivo* tRNA CRISPRi perturbations. **(A)** Heatmap showing *in vivo* tRNA CRISPRi perturbation profiles, with hierarchical clustering applied to group similar response patterns. **(B)** Scatter plot comparing the perturbation effects of aaRSs and tRNA-modifying enzymes estimated in this study with those reported in DepMap across all available cell lines. Each point represents one aaRS or tRNA-modifying enzyme. *R* and *P* values were calculated using Pearson correlation. **(C)** Bar plots showing the relative expression of target tRNA molecules upon single-guide CRISPRi-mediated tRNA knockdown in MDA-MB-231 cells compared to the non-targeting control (gCTR), quantified by RT–qPCR. Guide numbers in parentheses correspond to those used in the pooled CRISPRi screen. 5S rRNA served as the endogenous control for RT–qPCR quantification. Primers are listed. Bars represent the mean of biological and technical replicates, and error bars denote the standard error. *P* values were calculated from raw measurements using one-tailed *t* tests.

**Figure S2.**
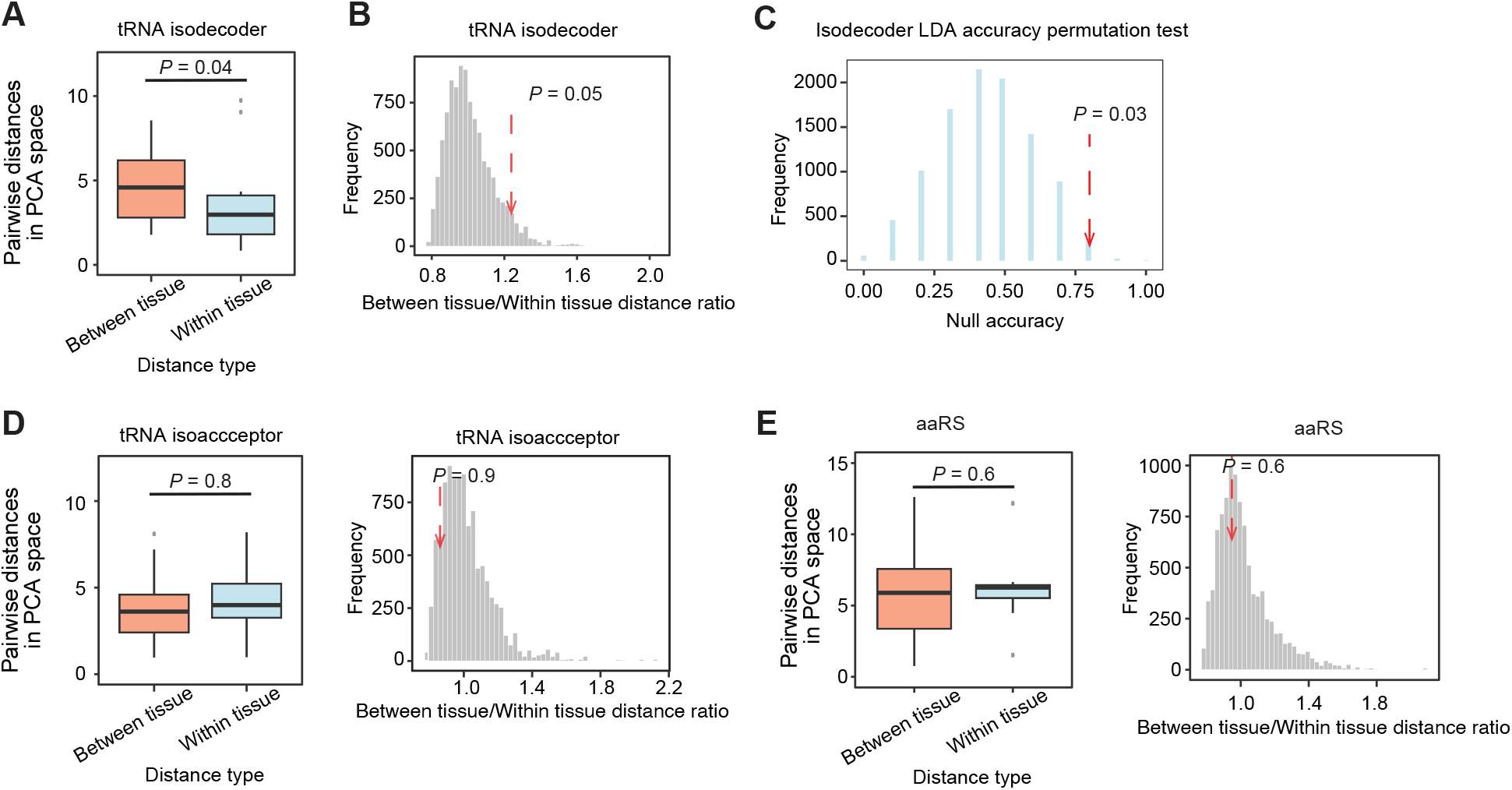
Analyses of tissue-specific CRISPRi perturbation profiles of tRNA isodecoders, isoacceptors, aaRSs, and tRNA-modifying enzymes. **(A)** Boxplots showing the distribution of pair-wise within-tissue and between-tissue Euclidean distances in PCA space (from Figure 1D) for tRNA isodecoder perturbation scores. Within-tissue distances were calculated only for tissues with at least three cell lines. Boxplots represent the median (center line), interquartile range (box), and 1.5 × interquartile range (whiskers); individual points indicate outliers. **(B)** Histogram showing the null distribution of the ratio between between-tissue and within-tissue Euclidean distances in PCA space (from Figure 1D) for tRNA isodecoder perturbation scores, obtained by random shuffling (*N* = 10,000). **(C)** Histogram showing the null distribution of LDA classifier accuracy in PCA space (from Figure 1D) for tRNA isodecoder perturbation scores, obtained by random shuffling (*N* = 10,000). **(D)** Similar to **(A–B)**, but performed at the tRNA isoacceptor level. **(E)** Similar to **(A–B)**, but performed for aaRS perturbation profiles.

We next asked whether these coordinated dependencies simply reflected shared decoding identity among related tRNAs. If this were the case, coordinated relationships would be expected to occur preferentially among isodecoders belonging to the same anticodon family. However, significant isodecoder coordination pairs were not enriched within the same anticodon family across breast, colon, or lung (**Figure S3F**; *P* = 1.0, 0.1, and 0.8, respectively). Thus, the coordinated organization of isodecoder dependencies cannot be explained simply by shared anticodon identity, suggesting that additional regulatory features contribute to these relationships.

Given our recent finding that tRNA transcription can be regulated in a gene-specific manner by chromatin-associated factors (Chen et al., 2025), we next asked whether coordinated isodecoder dependencies were associated with shared transcriptional regulation at their corresponding tDNA loci. H3K4me3 has been associated with active Pol III transcription through its relationship with a permissive chromatin environment (Oler et al., 2010; Barski et al., 2010). We therefore examined whether coordinated isodecoder dependencies were associated with shared chromatin states at their corresponding tDNA loci. Using ENCODE datasets (ENCODE Project Consortium, 2012) from tissue-matched cancer cell lines (MCF7 for breast, HCT116 for colon, and A549 for lung), we quantified H3K4me3 enrichment at tDNA loci corresponding to significant isodecoder pairs. Across all tissues, 236 of 426 (55.4%) gene-level pairs exhibited concurrent H3K4me3 enrichment (**Figure S3G**; 52.1% in breast, 31.7% in colon, and 69.5% in lung). Coordinated isodecoder pairs exhibited significantly greater concurrent H3K4me3 enrichment than non-coordinated isodecoder pairs (onesided empirical *P* = 0.04). Together, these results associate tissue-specific isodecoder co-dependencies with coordinated regulatory states at individual tDNA loci, supporting locus-specific transcriptional regulation as one potential contributor to their tissue-specific organization.

### 2.3 Isoacceptor dependencies couple codon-decoding demand to tumor physiology through an asymmetric tRNA-aaRS architecture

Unlike isodecoders, tRNA isoacceptor perturbations were largely tissue invariant, resembling the dependency patterns observed for aaRS-targeting perturbations (**Figure 1D-G**). We hypothesized that these tissue-invariant dependencies reflect fundamental translational requirements for sustaining cell proliferation under the shared fitness constraints imposed by *in vivo* tumor growth. We therefore sought to determine the biological principles underlying tRNA isoacceptor dependency.

We first asked whether isoacceptor dependencies could be explained by properties of the amino acids they decode. Pairwise correlation analysis revealed substantial similarity among isoacceptor dependency profiles across cancer cell lines (**Figure 2A**). However, neither codon-decoding redundancy nor the propensity of the encoded amino acid to undergo alternative decoding in human tumors (Tsour et al., 2026) explained the observed dependency structure (**Figure 2B**). Likewise, amino acid chemical class, molecular size, and other physicochemical characteristics showed little association with isoacceptor dependency (**Figure 2B**). Thus, the shared organization of isoacceptor dependencies is not primarily determined by intrinsic properties of the amino acids they decode.

**Figure 2.**
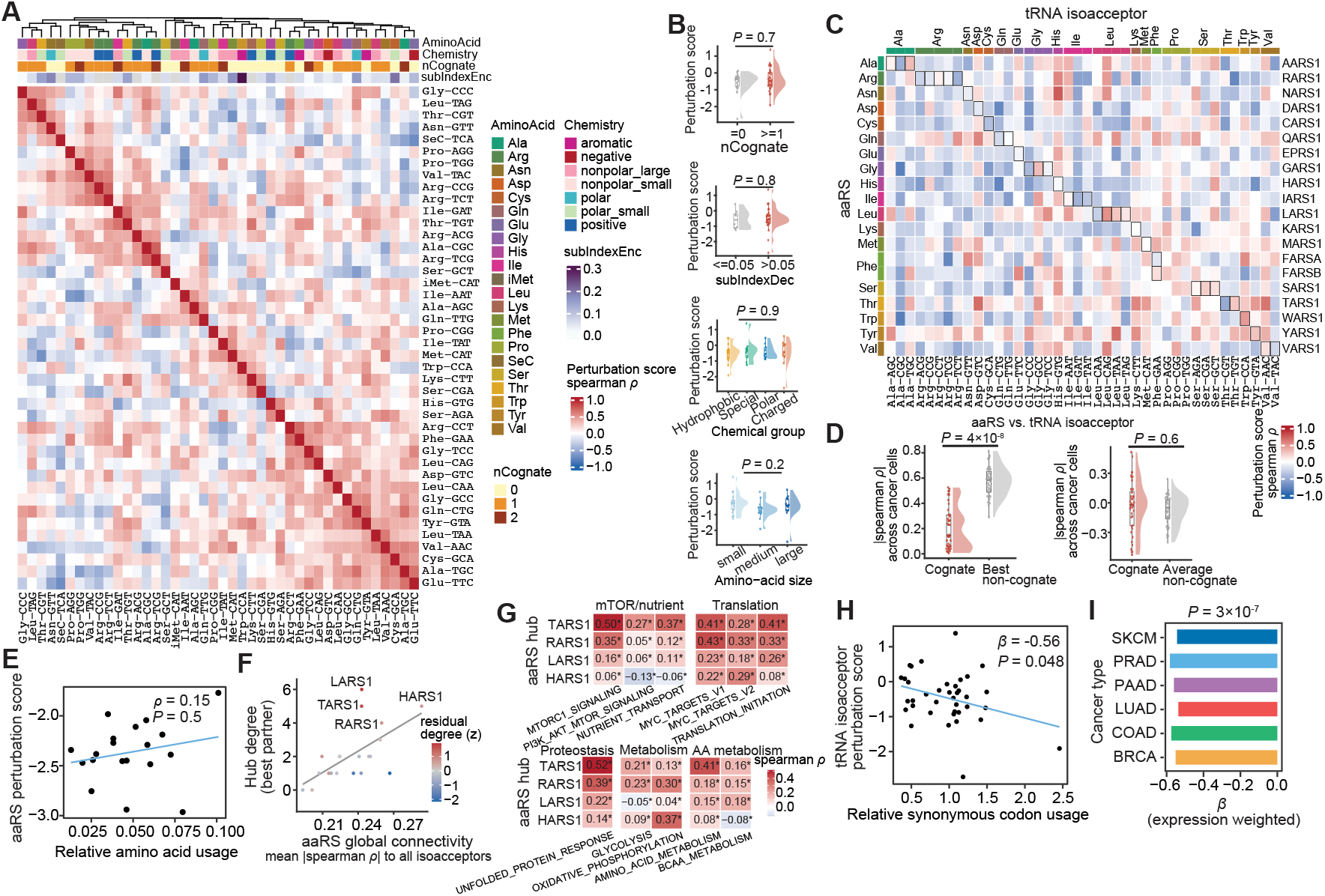
Functional organization of tRNA dependencies across tRNA isoacceptors and aminoacyl-tRNA synthetase levels *in vivo*. **(A)** Heatmap showing pairwise Spearman correlations between *in vivo* perturbation profiles of individual tRNA isoacceptors across cancer cell lines. Isoacceptors were hierarchically clustered according to perturbation similarity. Annotation tracks indicate the cognate amino acid, amino acid chemical class, codon-decoding redundancy, and the encoded amino acid substitution propensity derived from human tumor proteomic data. Codon-decoding redundancy (*nCognate*) represents the extent to which an isoacceptor shares decoded synonymous codons with other isoacceptors of the same amino acid. Encoded substitution propensity (*subIndexEnc*) summarizes the relative tendency of each encoded amino acid to undergo alternative decoding in human tumors. Lysine and arginine are not annotated because encoded substitutions cannot be reliably estimated from tryptic proteomic data. **(B)** Raincloud plots showing the distribution of average *in vivo* perturbation across tRNA isoacceptors grouped by amino acid chemical class, amino acid size, codon-decoding redundancy (*nCognate*), and encoded amino acid substitution propensity (*subIndexEnc*). Annotation categories are identical to those shown in **(A).** Average dependency was calculated as the mean *in vivo* perturbation score across cancer cell lines for each tRNA isoacceptor. Each point represents one tRNA isoacceptor; boxes indicate the median and interquartile range, and density envelopes show the distribution across isoacceptors. Group differences were assessed using the Mann–Whitney *U* test for two-group comparisons or the Kruskal–Wallis test for multi-group comparisons. **(C)** Heatmap showing Spearman correlations between *in vivo* perturbation profiles of individual aaRSs and individual tRNA isoacceptors across matched cancer cell lines. Rows represent aaRS genes and columns represent tRNA isoacceptors, ordered by decoded amino acid. Colored side bars indicate decoded amino acid identity. Outlined cells mark cognate aaRS–tRNA amino acid pairs. **(D)** Comparison of cognate and non-cognate aaRS associations for individual tRNA isoacceptors. For each tRNA isoacceptor, Spearman correlations between its *in vivo* perturbation profile and the perturbation profile of every aminoacyl-tRNA synthetase (aaRS) were calculated across matched cancer cell lines. Correlation strength was quantified as the absolute Spearman correlation (|ρ|). Left, comparison of the strongest cognate association (maximum |ρ| among aaRS charging the decoded amino acid) and the strongest non-cognate association (maximum |ρ| among all remaining aaRSes) for each isoacceptor. Right, comparison of the mean cognate and mean non-cognate association strengths for each isoacceptor. Each point represents one tRNA isoacceptor; box plots indicate the median and interquartile range. *P*-value was calcualted using paired Mann–Whitney *U* test tests. **(E)** Amino acid usage versus aaRS dependency. Amino acid usage was estimated from genome-wide coding sequences by summing codon frequencies for codons encoding the same amino acid and averaging across genes. For each aaRS, average *in vivo* dependency was calculated as the mean perturbation score across screened cancer cell lines. Points represent aaRS genes mapped to their cognate amino acid. ρ and *P* values were calculated with Spearman correlation. **(F)** Identification of recurrent aaRS hubs associated with tRNA isoacceptor dependencies. Global connectivity was defined as the mean absolute Spearman correlation between each aaRS and all tRNA isoacceptors across cancer cell lines. Hub degree was defined as the number of tRNA isoacceptors for which an aaRS exhibited the strongest dependency association (maximum absolute Spearman correlation). Each point represents one aaRS. Point color indicates the standardized residual from a linear model predicting hub degree from overall connectivity and dependency characteristics, with positive values indicating aaRSs exhibiting more hub connections than expected. **(G)** Association of aaRS hub expression with tumor physiological states across human cancers. Gene-set activity was quantified in TCGA PanCanAtlas primary tumors using single-sample gene set enrichment analysis (ssGSEA). Heatmap colors indicate the mean within-cancer-type Spearman correlation between aaRS expression and pathway activity across cancer types. Asterisks denote significant associations identified by a linear mixed-effects model with cancer type included as a random intercept (Benjamini–Hochberg-adjusted *P* < 0.05). **(H)** Relationship between isoacceptor dependency and synonymous codon usage. Relative synonymous codon usage (RSCU) was calculated from coding sequences by comparing the observed frequency of each codon with the expected frequency under equal usage of synonymous codons encoding the same amino acid. Tryptophan and methionine were excluded because each is encoded by a single codon. Each point represents one tRNA isoacceptor–codon pair. Isoacceptor dependency was defined as the average *in vivo* perturbation score across cancer cell lines. The association between RSCU and isoacceptor dependency was evaluated using a linear mixed-effects model accounting for amino acid identity. **(I)** Expression-weighted codon demand across tumor types. TCGA PanCanAtlas primary tumor RNA-seq profiles were used to estimate cancer-type-specific codon demand by weighting gene-level codon frequencies by mean gene expression within each cancer type. Codon demand was normalized within each amino acid to quantify relative synonymous codon demand. For each cancer type, tRNA isoacceptor dependency was averaged across matched cell lines and compared with expression-weighted synonymous codon demand. Bar heights show the estimated association between codon demand and isoacceptor dependency for each cancer type. Overall significance was assessed using a linear mixed-effects model accounting for amino acid identity and cancer type.

Because aaRSs directly charge their cognate tRNA isoacceptors, we next asked whether dependencies on these two components of the translational machinery were similarly organized. Surprisingly, the strongest aaRS–tRNA dependency associations frequently involved non-cognate rather than cognate pairs (**Figure 2C– D**), and cognate pairs were not significantly more correlated than the average non-cognate pair (**Figure 2D**). Thus, despite their direct biochemical relationship, aaRSs and their cognate tRNA isoacceptors do not exhibit correspondingly coupled fitness requirements, revealing an unexpected asymmetry in their dependency organization in the physiological tumor environment.

We next asked what biological principles might underlie this asymmetry by examining the organization of aaRS dependencies. Overall amino acid usage showed little association with aaRS dependency (**Figure 2E**), and amino acid chemical, physicochemical, and evolutionary properties likewise explained little of its variation (**Figure S4A–B**). Instead, several recurrent aaRS hubs, including LARS1 and TARS1, were associated with dependencies on multiple distinct tRNA isoacceptors (**Figure 2F**), suggesting that the observed aaRS dependencies during tumor growth might not be determined solely by their canonical aminoacylation relationships. Because aaRSs can also participate in nutrient sensing and signaling (Yoon et al., 2026), we asked whether their dependencies instead reflect broader aspects of tumor physiological state. Consistent with this possibility, recurrent aaRS hubs were preferentially associated with distinct physiological programs in human tumors. For example, LARS1 and TARS1 were associated with nutrient transport, amino acid metabolism, mTOR signaling, and translational programs (**Figure 2G**). Moreover, aaRS dependencies exhibited distinct amino acid-specific relationships with estimated amino acid flux derived from TCGA transcriptomes (**Figure S4C–D**) (Huang et al., 2023; Cancer Genome Atlas Research Network et al., 2013). Together, these observations suggest that aaRS dependencies are shaped not only by translational decoding but also by broader tumor physiological states.

Having established that neither amino acid properties nor cognate aaRS relationships explain isoacceptor dependency, we next asked whether the primary organizing principle instead resides at the level of codon decoding. Genome-wide synonymous codon usage was strongly associated with isoacceptor dependency, such that frequently decoded synonymous codons were associated with stronger dependency (more negative perturbation scores) on their cognate tRNA isoacceptors (**Figure 2H**). Moreover, expression-weighted codon demand derived from TCGA transcriptomes showed consistent associations with isoacceptor dependency across multiple tumor types (**Figure 2I**), indicating that physiology-specific translational demand further shapes these decoding requirements under *in vivo* conditions. Together, these results reveal distinct organizational principles governing the two arms of the translation machinery. Unlike aaRS dependencies, which reflect both translational demand and broader amino acid-specific metabolic and physiological states, tRNA isoacceptor dependencies are fundamentally organized by codon decoding demand.

### 2.4. Parallel *in vitro* cell culture extensively reorganizes the tRNA dependency landscape

Our *in vivo* screens revealed a highly structured tRNA dependency landscape, with tissue-specific organization emerging at the isodecoder level (**Figure S3**) and isoacceptor dependencies associated with codon-decoding demand (**Figure 2H–I**) during physiological tumor growth. However, because these dependencies were measured in established tumors, they could reflect both intrinsic properties of the cancer cells and requirements imposed by the tumor environment. We therefore asked whether this dependency architecture is maintained when the same cancer cells are removed from the tumor environment. Using the same cancer cell lines and perturbation library, we performed matched *in vitro* cell culture CRISPRi screens in parallel with our *in vivo* screens (**Figures 3A** and **S5A**). Dependencies on aaRSs and tRNA-modifying enzymes remained substantially concordant between *in vitro* and *in vivo* growth (**Figures 3B** and **S5B**), indicating that a large component of these dependencies is preserved across environments. This concordance also provided an internal benchmark demonstrating that the *in vitro* screens retained sufficient biological signal to recover reproducible dependencies across conditions. In marked contrast, individual tRNA dependencies showed substantially weaker concordance between the two conditions (**Figure 3B** and **S5C**), suggesting that the tRNA dependency landscape is considerably more sensitive to growth environment.

**Figure 3.**
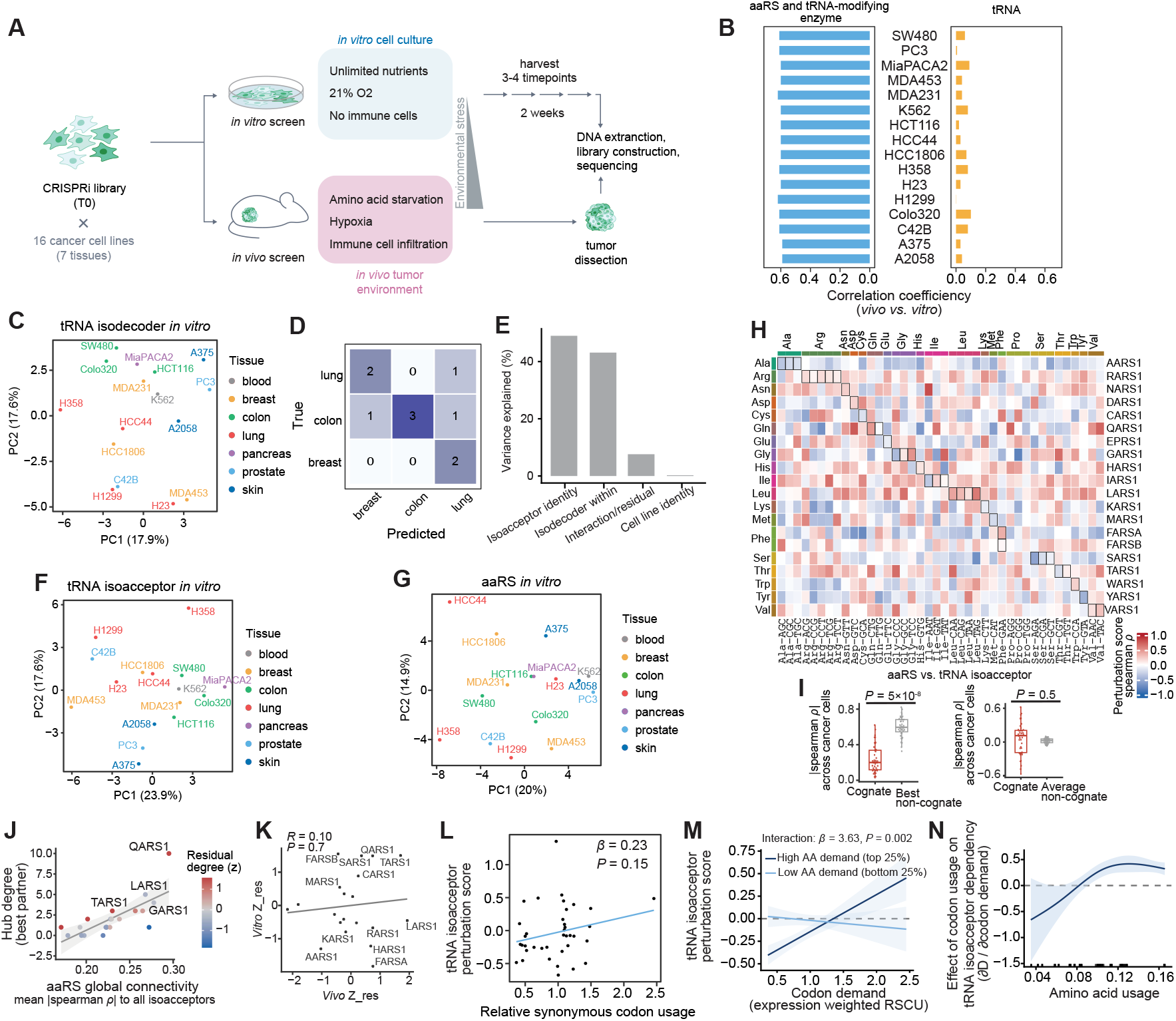
Reorganized tRNA isodecoder and isoacceptor dependency landscapes across parallel *in vitro* CRISPRi screens. **(A)** Schematic of parallel *in vivo* and *in vitro* tRNA CRISPRi screens across cancer cell lines. **(B)** Pearson correlation coefficients of perturbation effects between *in vivo* and *in vitro* screens for tRNA molecules (orange) and protein-coding genes, including aaRSs and tRNA-modifying enzymes (blue). **(C)** Principal component analysis (PCA) of tRNA isodecoder perturbation profiles across cancer cell lines cultured *in vitro*, colored by tissue of origin. Analysis is analogous to Fig. 1D. **(D)** Confusion matrix of leave-one-out (LOO) supervised classification of tissue of origin from *in vitro* tRNA isodecoder perturbation profiles. Analysis is analogous to Fig. 1E. **(E)** Variance decomposition of *in vitro* isodecoder dependencies. Isodecoder dependency scores across cell lines were partitioned into variation attributable to isoacceptor identity, isodecoder identity nested within isoacceptor, cell line identity, and the remaining isodecoder-by-cell-line interaction/residual. Bars show the percentage of total sum of squares (% SS) attributable to each component. **(F–G)** PCA of *in vitro* perturbation profiles at the tRNA isoacceptor level (F) and for aaRS genes (G), similar to (C). **(H)** Heatmap of Spearman correlations between *in vitro* perturbation profiles of individual aaRSs and tRNA isoacceptors across matched cancer cell lines. Rows represent aaRS genes and columns represent tRNA isoacceptors, ordered by decoded amino acid. Side bars indicate amino acid identity, and outlined cells denote cognate aaRS–tRNA amino acid pairs. Analysis is analogous to Fig. 2C. **(I)** Comparison of cognate and non-cognate aaRS associations for individual tRNA isoacceptors. Left, strongest cognate association (maximum |ρ| among aaRSs charging the corresponding amino acid) versus strongest non-cognate association (maximum |ρ| among all other aaRSs). Right, mean cognate versus mean non-cognate association strength. Each point represents one tRNA isoacceptor; box plots indicate the median and interquartile range. *P* values were calculated using paired Mann–Whitney *U* tests. Analysis is analogous to Fig. 2D. **(J)** Identification of recurrent aaRS hubs associated with *in vitro* tRNA isoacceptor dependencies. Point color indicates standardized residual hub degree from a linear model accounting for overall aaRS connectivity and dependency characteristics; positive values indicate aaRSs with more hub connections than expected. The gray line shows the linear regression of hub degree on global aaRS connectivity, with the shaded region indicating the 95% confidence interval. Analysis is analogous to Fig. 2F. **(K)** Comparison of standardized residual hub degree between *in vitro* and *in vivo* conditions. Each point represents one aaRS. *In vivo* standardized residual hub degree is defined as in Fig. 2F, and *in vitro* standardized residual hub degree as in (J).

**Figure S4.**
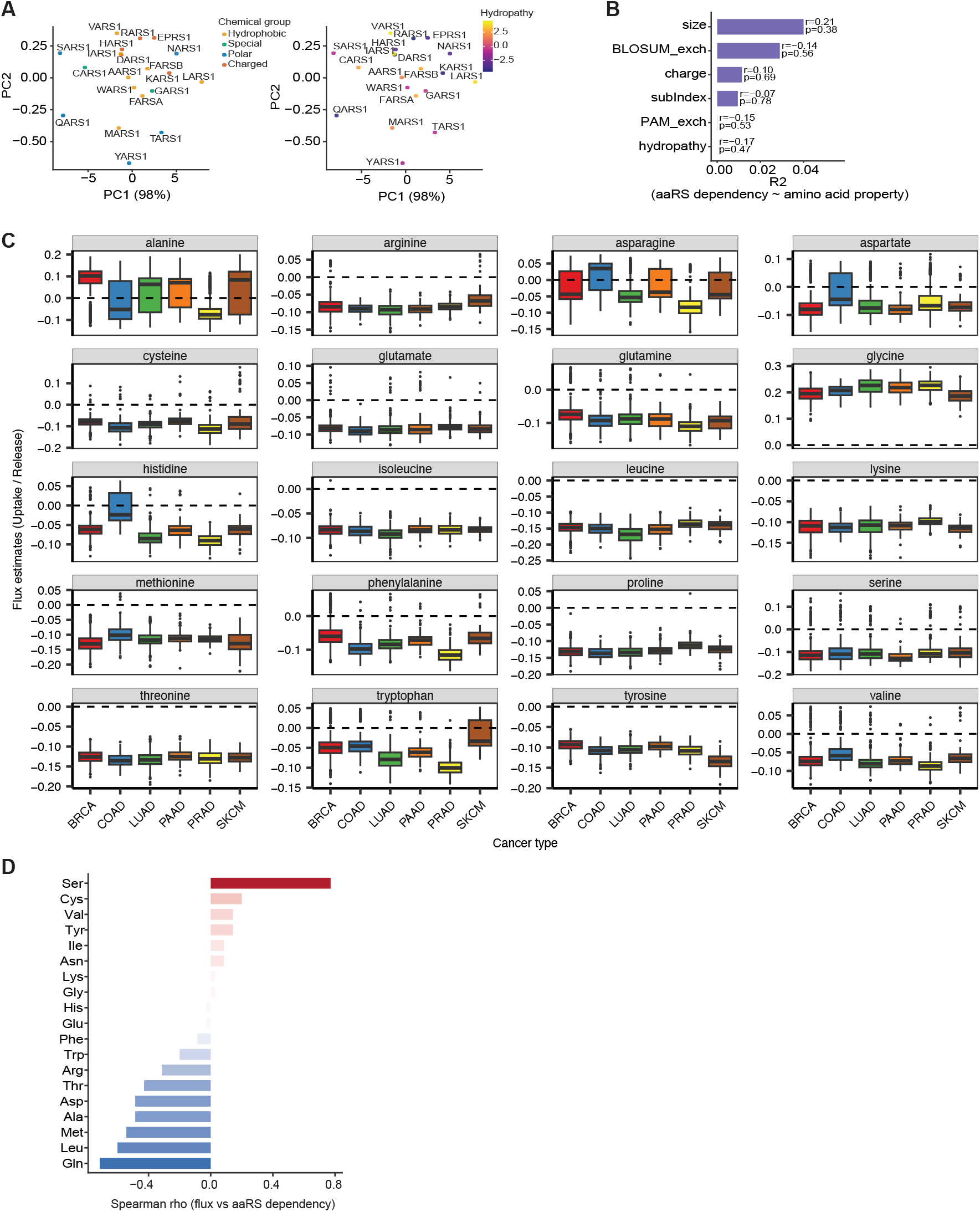
aaRS dependencies in tumor physiological states. **(A)** Principal component analysis (PCA) of aminoacyl-tRNA synthetase (aaRS) dependency profiles across cancer cell lines. The same projection is colored by amino-acid chemical class (left) and hydropathy (right). **(B)** Variance in mean aaRS dependency explained (*R*^2^) by individual amino-acid properties. BLOSUM_exch and PAM_exch represent the mean off-diagonal substitution scores derived from the BLOSUM62 and PAM250 matrices, respectively. subIndexEnc denotes the encoded amino acid substitution propensity. **(C)** Inferred amino acid metabolic flux across TCGA cancer types. Amino acid exchange fluxes were inferred from TCGA RNA-seq data. **(D)** Amino-acid-specific associations between metabolic flux and aaRS dependency. For each amino acid, Spearman correlations were calculated between median metabolic flux and mean dependency of the cognate aaRS across matched cancer types. aaRS dependency was averaged across cell lines within each TCGA-matched lineage and assigned to the amino acid charged by each synthetase before integration with metabolic flux estimates.

To determine how this environmental sensitivity reshapes tRNA dependency, we first characterized the *in vitro* landscape using the same hierarchical framework applied *in vivo*. At the isodecoder level, the clear tissue-associated organization observed during tumor growth was largely absent under *in vitro* conditions (**Figures 3C** and **S6A-B**). We therefore asked whether this loss reflected a general attenuation of tissue separation or, more specifically, disruption of the isodecoder relationships that distinguished tissues *in vivo*. To distinguish these possibilities, we directly followed the tissue-specific isodecoder pairs identified from the *in vivo* screens (**Figure S3A–C**) and quantified their tissue-specific coherence *in vivo* and *in vitro* by comparing similarity among cell lines within the corresponding tissue with similarity across tissues (**Figure S6C**). Across breast, colon, and lung cancer cells, these relationships exhibited substantially stronger within-tissue coherence *in vivo* than *in vitro*, whereas the corresponding tissue organization was weak or absent in culture (**Figure S6D-F**). Thus, removal from the tumor environment is accompanied by attenuation of the specific isodecoder relationships that organize tRNA dependencies within tissues. Our early observation of the persistence of transcriptional coordination among these *in vivo*-defined pairs in ENCODE cell-line data (**Figure S3G**) nevertheless suggests that their underlying regulatory relationships are not entirely lost in culture, but that their functional manifestation is affected by the physiological tumor environment.

The attenuation of tissue organization raised the question of what instead structures isodecoder dependencies under *in vitro* conditions. We therefore decomposed isodecoder dependency variation into contributions from isoacceptor identity, isodecoder identity nested within isoacceptor, cell-line identity, and the remaining isodecoder-by-cell-line interaction/residual (**Figure 3E**). Isoacceptor identity and isodecoder identity within isoacceptors accounted for 49.0% and 43.1% of the total sum of squares, respectively, whereas cell-line identity accounted for only 0.25%. It indicates that the loss of tissue-associated organization did not render the *in vitro* dependency landscape unstructured. Instead, dependency variation remained strongly organized by tRNA identity, with substantial contributions from both isoacceptor and isodecoder levels.

We next asked whether environmental reorganization extended to the relationship between isoacceptor and aaRS dependencies. Similar to our observations *in vivo*, principal component analysis of both isoacceptor and aaRS dependencies showed little tissue-associated separation under *in vitro* conditions (**Figures 3F-G** and **3G-H**), and the asymmetric relationship between aaRS and isoacceptor dependencies remained evident in *in vitro* cell culture (**Figure 3H-I**). However, the aaRS hubs underlying this architecture differed markedly between environments (**Figure 3J**). aaRSs exhibiting higher-than-expected connectivity *in vitro* were distinct from the recurrent hubs identified *in vivo*, and aaRS hub scores were uncorrelated between conditions (*R* = 0.1, *P* = 0.7; **Figure 3K**). Notably, LARS1, a recurrent hub *in vivo* associated with tumor physiological programs (**Figure 2F-G**), did not exhibit significantly higher-than-expected connectivity *in vitro* (**Figure 3J**). Together, aaRS–isoacceptor asymmetry is preserved across growth environments, whereas the specific aaRS relationships underlying this architecture are environmentally reorganized.

Finally, because codon-decoding demand was a major organizing principle of isoacceptor dependency during physiological tumor growth, we asked whether the same relationship was preserved in cell culture. Relative synonymous codon usage alone was not significantly associated with isoacceptor dependency *in vitro* (**Figure 3L**), indicating that the direct relationship observed during tumor growth was not maintained under culture conditions. However, this did not represent a complete loss of codon-dependent organization. Instead, the relationship between synonymous codon demand and isoacceptor dependency changed with amino acid demand. Among amino acids with low demand, increasing synonymous codon demand was associated with more negative perturbation scores, consistent with the relationship observed during physiological tumor growth. This relationship shifted among amino acids with high demand (**Figure 3M**; interaction *P* = 0.002). Modeling amino acid demand continuously further revealed a progressive change in the effect of synonymous codon demand, from negative at low amino acid demand to positive at high amino acid demand (**Figure 3N**). Thus, the lack of an overall codon-demand association *in vitro* masks distinct relationships across the amino acid-demand spectrum: the expected association between codon demand and tRNA requirement is retained at low amino acid demand but progressively shifts as amino acid demand increases. Together, these results indicate that the relationship between translational demand and tRNA requirement is fundamentally reorganized in cell culture.

**Figure S5.**
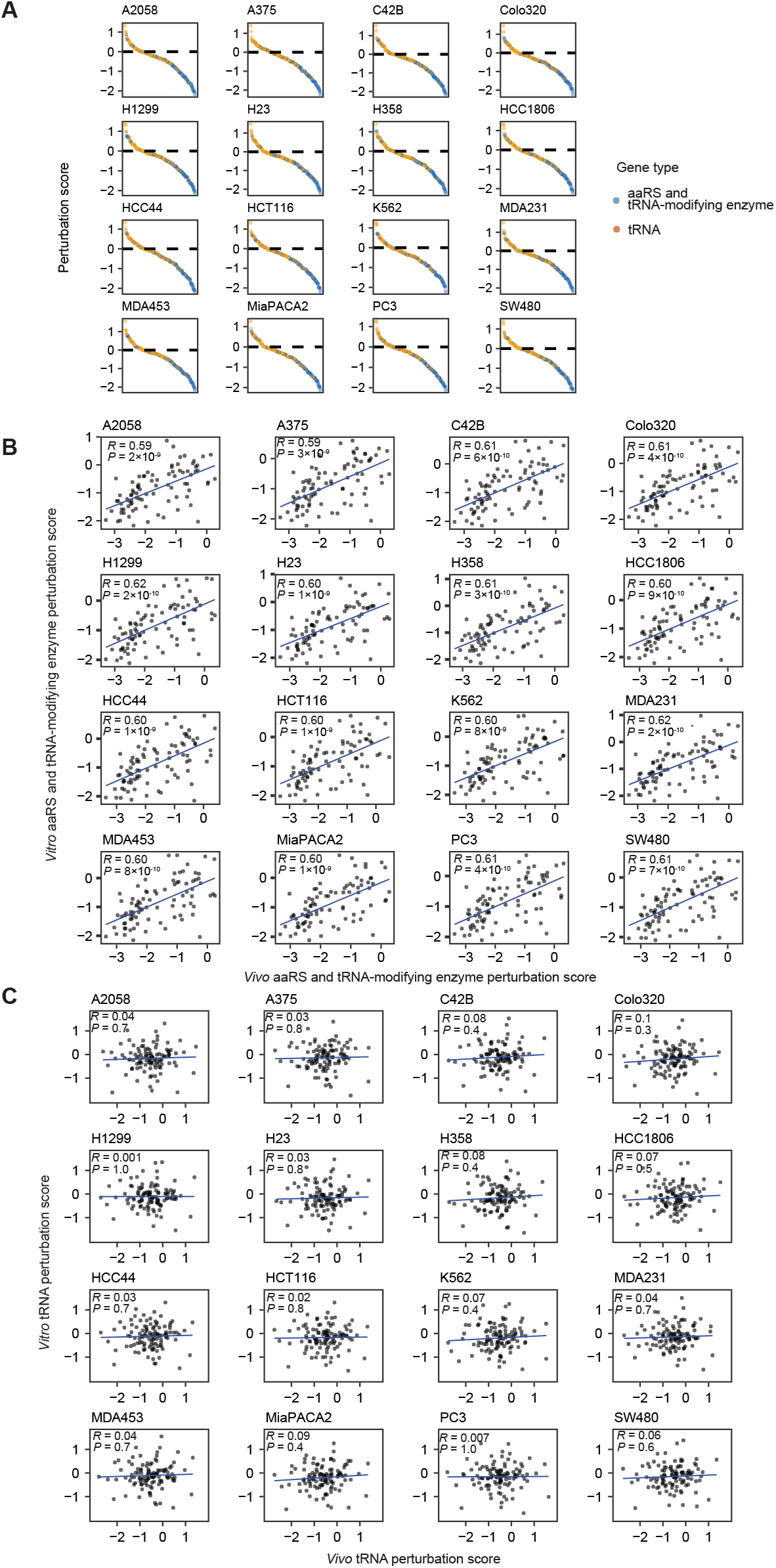
Comparison of *in vivo* and *in vitro* CRISPRi perturbation effects. **(A)** Waterfall plots showing the distribution of perturbation effects for tRNA molecules, aminoacyl-tRNA synthetases (aaRSs), and tRNA-modifying enzymes across individual cell lines. **(B)** Scatter plot comparing the perturbation effects of aaRSs and tRNA-modifying enzymes between *in vivo* and *in vitro* CRISPRi experiments. Pearson correlation coefficients (*R*) and corresponding *P* values are shown. **(C)** Scatter plot comparing tRNA perturbation effects between *in vivo* and *in vitro* CRISPRi experiments. Pearson correlation coefficients (*R*) and corresponding *P* values are shown.

Figure 3 | **(L)** Relationship between *in vitro* isoacceptor dependency and synonymous codon usage. Relative synonymous codon usage (RSCU) was calculated as in Fig. 2H. The association between RSCU and isoacceptor dependency was evaluated using a linear mixed-effects model accounting for amino acid identity. **(M)** Relationship between expression-weighted codon demand and *in vitro* isoacceptor dependency at different levels of amino-acid demand. For each cell line, codon demand was calculated by weighting gene-specific codon frequencies by CCLE gene expression [log_2_ (TPM + 1)] and summing across expressed genes. Expression-weighted codon demand was normalized within each amino acid to quantify relative demand among synonymous codons. Predicted isoacceptor dependency is shown across codon demand at low (bottom 25%) and high (top 25%) amino-acid demand. Lines and shaded regions indicate model predictions and 95% confidence intervals, respectively. **(N)** Conditional effect of codon demand across the amino-acid demand spectrum. The estimated effect of expression-weighted codon demand on isoacceptor dependency (∂*D*/ ∂codon demand) is shown as a continuous function of amino-acid demand share. Estimates were obtained using a varying-coefficient generalized additive model including a smooth amino-acid-demand effect, an amino-acid-demand-dependent codon-demand effect, and cell line as a random effect. The solid line and shaded region indicate the estimated conditional effect and 95% confidence interval, respectively; the dashed horizontal line denotes no codon-demand association, and the rug along the x axis indicates the distribution of observed amino-acid demand values.

**Figure S6.**
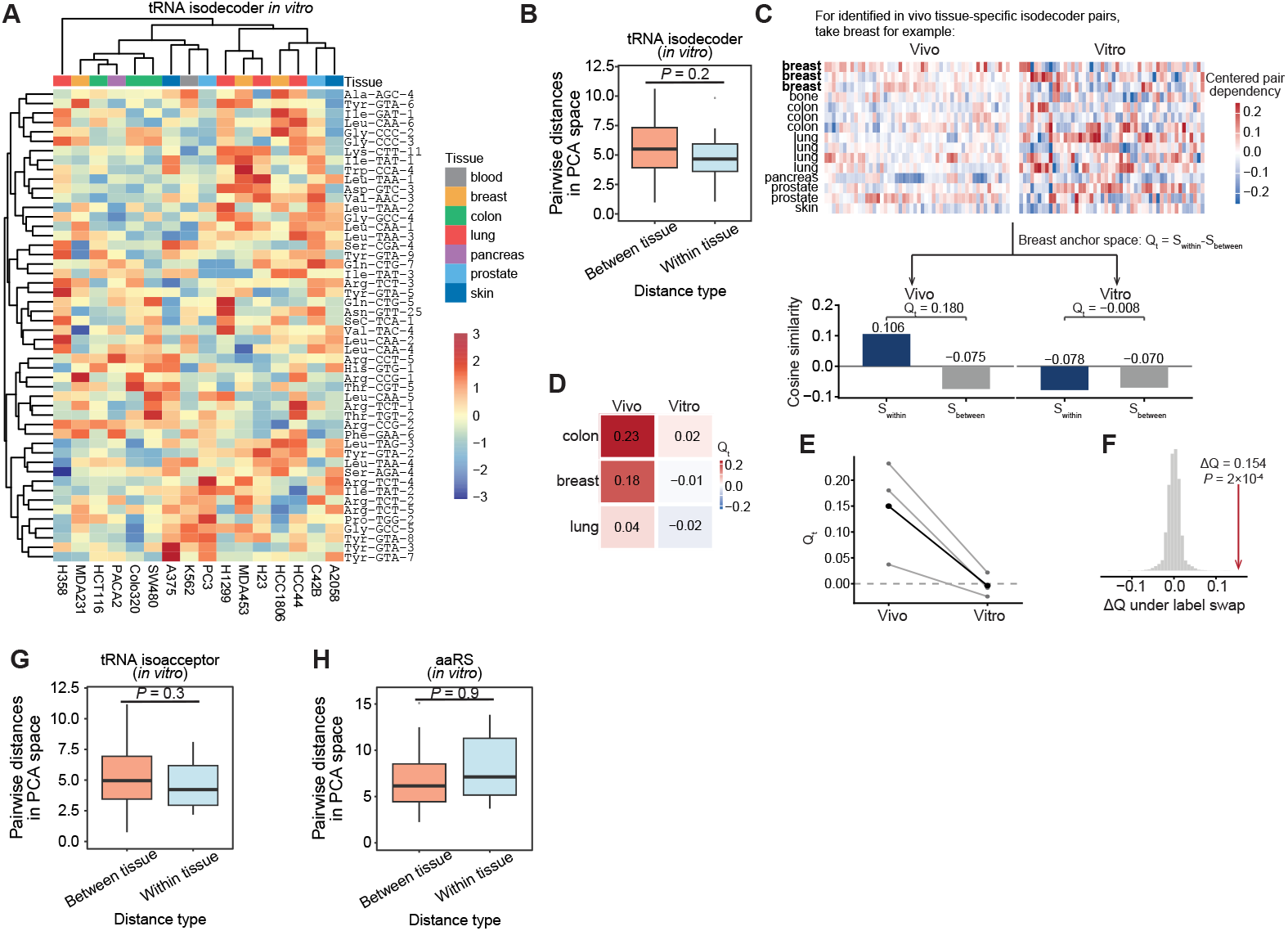
Organization of tRNA dependencies *in vitro*. **(A)** Heatmap of *in vitro* tRNA isodecoder CRISPRi perturbation profiles across cell lines. Isodecoders and cell lines were hierarchically clustered using Spearman-correlation-based distances to group similar dependency patterns. **(B)** Pairwise Euclidean distances between cell lines in the PCA space shown in (**Fig. 3C**). Within-tissue distances were calculated between cell lines from the same tissue, whereas between-tissue distances were calculated between cell lines from different tissues. Within-tissue comparisons were restricted to tissues represented by at least three cell lines. Boxplots show the median, interquartile range, and 1.5 × interquartile range; points indicate outliers. **(C)** Schematic and representative calculation of tissue-specific coherence using breast as the anchor tissue. Tissue-specific isodecoder pairs were identified from the *in vivo* dependency data (**Fig. S3**), and the same fixed pair set was evaluated under both *in vivo* and *in vitro* conditions. All 50 breast-specific isodecoder pairs are shown. **(D)** Tissue-specific coherence (*Q*_*t*_) for each anchor tissue under *in vivo* and *in vitro* conditions. Tissues are ordered by the magnitude of the *in vivo*-to-*in vitro* change in coherence. All three tissues showed reduced coherence *in vitro*, with values approaching zero. **(E)** Paired comparison of tissue-specific coherence between *in vivo* and *in vitro* conditions. Gray lines denote individual anchor tissues, and the black line denotes the equally weighted mean across tissues. **(F)** Permutation test for the observed change in tissue-specific coherence. *In vivo* and *in vitro* condition labels were swapped at the cell-line level while retaining the tissue-specific pair sets, and pooled Δ*Q* was recalculated for each permutation. The inset shows the resulting null distribution, with the observed Δ*Q* indicated in red. **(G)** Pairwise within-tissue and between-tissue Euclidean distances in the *in vitro* isoacceptor PCA space, calculated as in **(B). (H)** Pairwise within-tissue and between-tissue Euclidean distances in the *in vitro* aaRS dependency PCA space, calculated as in **(B)**.

### 2.5. Tumor growth induces a shared transition in tRNA isoacceptor dependency state

The extensive reorganization of tRNA dependencies between tumor growth and cell culture raised the question of whether these environmental differences reflect heterogeneous responses specific to individual cancer cells or a common transition superimposed on their distinct dependency landscapes. Because the same cell lines retain substantial differences in tRNA dependency across environments, however, the observed landscapes jointly encode stable features associated with cellular identity and changes associated with growth environment. We therefore sought to disentangle these intertwined sources of variation, separating dependency features that remain coupled to cellular identity (“who”) from those that are remodeled with growth environment (“where”).

To accomplish this, we developed a paired conditional variational autoencoder (pcVAE) that jointly models matched isoacceptor dependency profiles from the same cancer cell lines in the two growth environments (**Figure 4A**). The model represents each profile through an identity latent (*z*_id_), constrained to remain shared between matched conditions, and a state latent (*z*_state_), which captures dependency variation associated with growth environment (**Figure S7A**). The difference in state latents between matched *in vivo* and *in vitro* samples (Δ*z*_state_) therefore provides a representation of environment-associated dependency remodeling after separating variation associated with stable cellular identity. Model validation confirmed effective separation of these components: identity and state representations were effectively orthogonal (**Figure S7**B), matched samples retained highly concordant identity coordinates (**Figures 4B** and **S7C**), and swapping identity representations between cell lines substantially impaired reconstruction (**Figure S7D**). Thus, the pcVAE provides a framework for resolving stable and environmentally plastic components of tRNA isoacceptor dependency.

**Figure 4.**
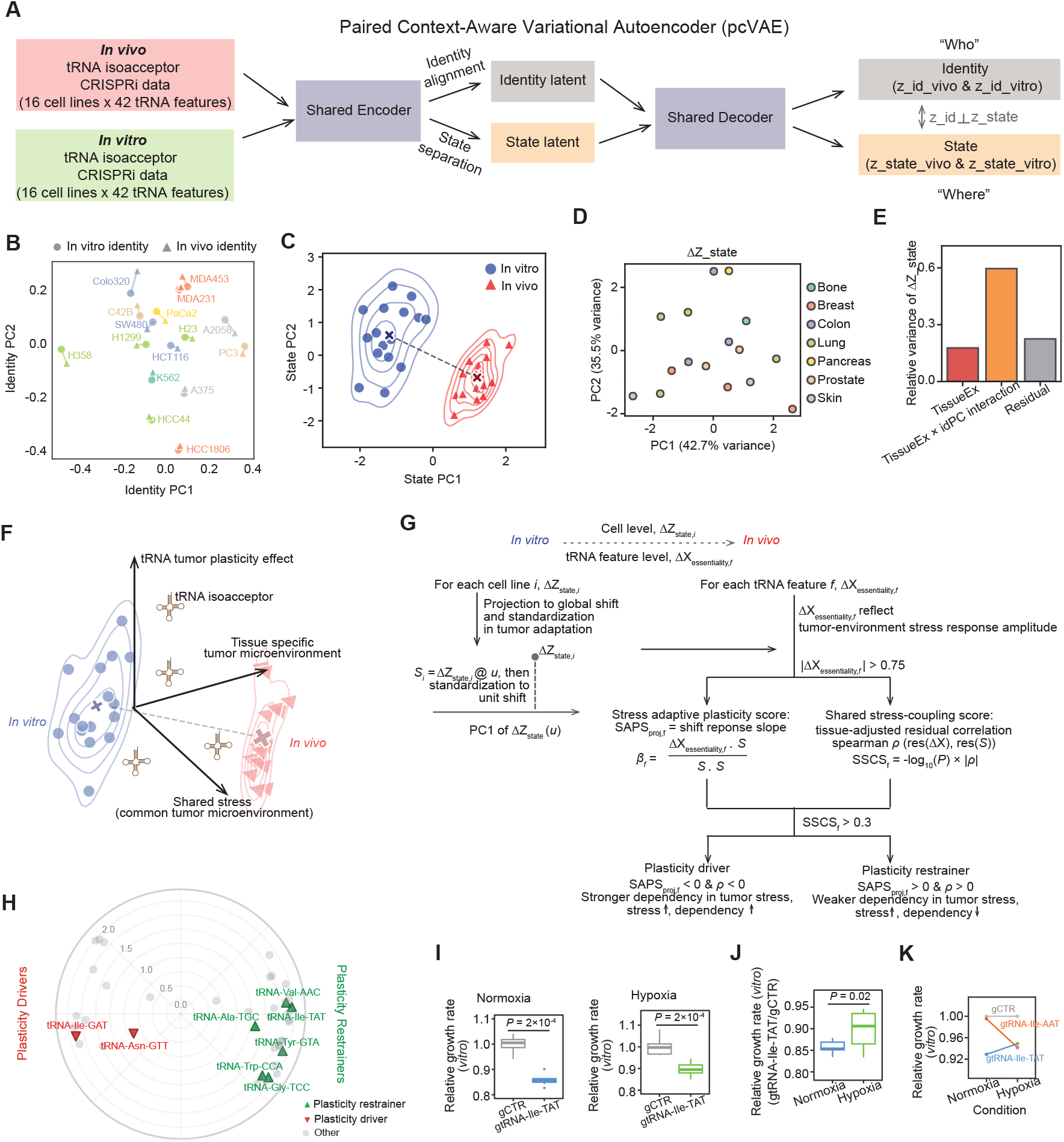
Disentangling cellular identity and environmental state reveals distinct modes of tRNA isoacceptor dependency plasticity. **(A)** Schematic of the paired context-aware variational autoencoder (pcVAE) framework used to disentangle cellular identity and adaptive state from paired *in vivo*–*in vitro* tRNA isoacceptor perturbation profiles. **(B)** PCA visualization of identity latent variables (*z*_id_) inferred by pcVAE for *in vivo* and *in vitro* cell lines. **(C)** Two-dimensional projection of cellular state transitions between *in vitro* and *in vivo* contexts. State PC1 and State PC2 represent the first two principal components of the combined *in vitro* and *in vivo* (*z*_state_) embeddings. **(D)** PCA of Δ*z*_state_ (*in vivo* – *in vitro*) for each cell line, colored by tissue type. **(E)** Variance decomposition of Δ*z*_state_ into tissue-extrinsic effects, interaction between identity and tissue-extrinsic components, and residual variance. Only tissues with at least three cell lines were included; bars show the mean across tissues. **(F)** Conceptual schematic illustrating how tumor-associated stress reshapes tRNA essentiality through shared and tissue-specific microenvironmental pressures. **(G)** Overview of the framework for classifying tRNA plasticity drivers and restrainers. For each tRNA isoacceptor feature, the Stress-Adaptive Plasticity Score (SAPS) quantifies the slope of the adaptive response (β _*f*_) from *state*_*in vitro*_ to *state*_*in vivo*_, standardized along a global tumor adaptation axis. In parallel, the shared stress-coupling score (SSCS) is computed using tissue-adjusted residual Spearman correlation to quantify monotonic coupling between tRNA isoacceptor dependency and stress progression. **(H)** Polar plot showing tRNA isoacceptors classified as plasticity drivers (red) or plasticity restrainers (green) defined from **(G)**.

We next examined how growth environment was represented after this disentanglement. Whereas matched samples remained closely aligned in identity space, their state latents separated strongly between *in vitro* and *in vivo* conditions (**Figures 4C** and **S7E-F**; separation ratio = 1.7, *P* < 0.001, permutation test). This state-latent separation suggests that tRNA isoacceptor dependencies undergo an environment-associated response, pointing to their potential plastic role in adaptation to the tumor environment. Importantly, we observed that the state-latent changes from *in vitro* to *in vivo* (**Figure 4D**) followed a highly consistent directional alignment across cancer cells (the length of the mean normalized flow vector, *R* = 0.8; **Figure S7G**), although heterogeneous alterations can observe after removing this global consistent component (*R* = 0.05; **Figure S7H**). These results indicate that despite substantial differences in baseline tRNA dependency among cancer cells in environmental response, physiological tumor condition induces a highly coherent transition in tRNA isoacceptor dependency state.

Decomposition of the state-latent changes further revealed contributions from both tissue context and cell-by-environment interactions, with tissue-associated effects explaining approximately 25% of the variation and cell-by-environment interactions accounting for more than 50% (**Figures 4E** and **S7I**). Together, these results reveal a structured plastic component of tRNA isoacceptor dependency, in which physiological tumor growth is associated with a broadly shared shift across cancer cells while cellular context influences how that shift is manifested.

### 2.6. tRNA isoacceptors define distinct modes of plasticity during adaptation to the tumor environment

The identification of a dominant axis of tRNA dependency remodeling between cell culture and physiological tumor growth revealed a common dimension of environmental dependency plasticity. The strong alignment of this response across cancer cells, together with the distinct responses of individual isoacceptors, suggested that different tRNAs may be differentially organized along this shared environmental axis. We next sought to determine how individual tRNA isoacceptor dependencies relate to progression along this response to the physiological tumor environment (**Figure 4F**).

Figure 4 | **(I)** Box plots showing proliferation rates following knockdown of a representative plasticity restrainer (gtRNA-Ile-TAT) compared with gCTR under normoxic (left) and hypoxic *in vitro* (right) conditions. Each condition includes eight replicates. *P* values were calculated using the Mann–Whitney *U* test. Boxes represent the interquartile range (IQR) with the median indicated; whiskers extend to 1.5 × IQR. **(J)** Box plot showing relative proliferation rates of gtRNA-Ile-TAT under normoxic and hypoxic *in vitro* conditions, derived from **(K)**. *P* value was calculated using a one-sided Mann–Whitney *U* test. Boxes represent the interquartile range (IQR) with the median indicated; whiskers extend to 1.5 × IQR. **(K)** Interaction plot showing relative *in vitro* growth rate shifts from normal to hypoxic conditions, illustrating the adaptive grammar captured by SAPS.

**Figure S7.**
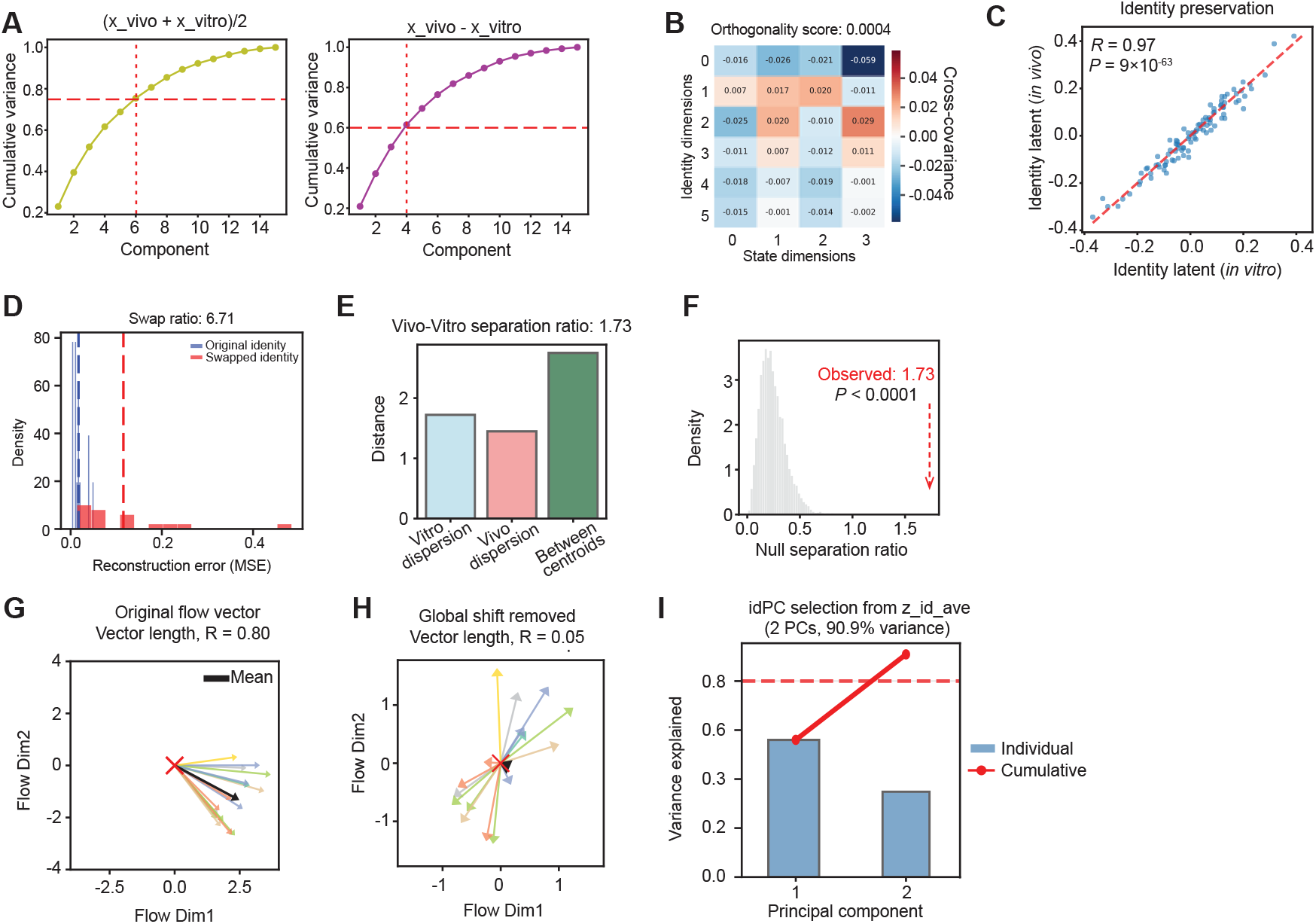
Quality control of the pcVAE model. **(A)** PCA variance explained for vivo vitro, and vivo vitro 2, used to guide the determination of the number of latent dimensions representing state and identity in the pcVAE model. **(B)** Heatmap showing the cross-covariance between identity latents and state latents. **(C)** Scatter plot comparing *in vivo* and *in vitro* identity components derived from the pcVAE model. Pearson correlation coefficients (*R*) and corresponding *P* values are shown. **(D)** Histogram showing the reconstruction error distribution for original versus swapped identity representations. **(E** Bar plot showing the dispersion of *in vivo* and *in vitro* state representations and the separation between their centroids, as presented in Figure 4C. **(F)** Histogram showing the null distribution of the *in vivo*–*in vitro* separation ratio shown in **(F)** and Figure 4C, generated from 10,000 random shuffles. **(G)** Vector-field representation of the directional flow from *in vitro* to *in vivo* states, with all vectors anchored at a common origin. Directional coherence was quantified by the vector-strength metric *R*, defined as the length of the mean normalized flow vector, where *R* = 1 indicates perfect directional alignment and *R* = 0 indicates no net directional coherence. **(H)** Flow field after removal of the global adaptation component, highlighting tissue-or cell-dependent residual dynamics. **(I)** Bar plot showing the variance explained by each principal component of *z*_id_ave_, with the red line indicating cumulative variance explained.

To quantify this relationship, we developed the Stress-Adaptive Plasticity Score (SAPS), which measures how the change in dependency of each isoacceptor scales with position along the shared environmental response (**Figure 4G**). We additionally assessed the consistency of this relationship across cellular contexts using a tissue-adjusted correlation between dependency remodeling and the shared response. These complementary measurements distinguish isoacceptors whose dependencies change consistently with increasing environmental remodeling from those showing the opposite relationship along this shared physiological tumor environment-associated axis.

This analysis revealed distinct classes of isoacceptors associated with the organization of physiological tumor environment-dependent tRNA plasticity (**Figure 4H**). We termed isoacceptors whose dependency behavior was positively associated with progression along the environmental response *plasticity drivers*, whereas isoacceptors exhibiting the opposing relationship were classified as *plasticity restrainers*. Importantly, these classifications describe how individual tRNA dependencies are organized relative to the physiological tumor environmental response. Therefore, our pcVAE-SAPS framework identifies specific tRNA isoacceptors whose dependencies systematically distinguish alternative modes of plasticity during the transition from cell culture to physiological tumor growth.

These opposing modes of plasticity during adaptation to the tumor environment were distributed across distinct amino-acid families rather than being restricted to a particular decoding group. For example, tRNA-Ile-GAT and tRNA-Asn-GTT exhibited driver-associated behavior, whereas tRNA-Val-AAC, tRNA-Ile-TAT, and tRNA-Gly-TCC showed the opposing pattern (**Figure 4H**). Thus, physiological tumor growth does not uniformly increase or decrease tRNA requirements, nor does it selectively remodel a single amino-acid family; instead, individual isoacceptors distributed across the decoding repertoire exhibit distinct directions of dependency remodeling along the same environmental response. Notably, tRNAs decoding the same amino acid could also exhibit opposing modes of plasticity: tRNA-Ile-GAT showed driver-associated behavior, whereas tRNA-Ile-TAT showed a restrainer-associated pattern. Because these isoacceptors supply the same amino acid but decode distinct synonymous codons, their opposing behavior suggests that environmental tRNA dependency plasticity may be organized at the level of codon-specific decoding demand. Together, these observations suggest that codon-specific decoding may represent an important organizing principle of tRNA dependency plasticity during adaptation to the tumor environment.

### 2.7. Plasticity-associated tRNA isoacceptors show differential dependencies under tumor-associated stress

The identification of plasticity drivers and restrainers from matched *in vivo* and *in vitro* screens raised the question of whether their predicted dependency behavior could be reproduced under a defined environmental pressure. Although the physiological tumor environment comprises multiple interacting environmental factors, testing individual pressures provides an opportunity to determine whether specific components of this complex environment are sufficient to alter tRNA dependency. We therefore examined hypoxia, a common feature of the tumor microenvironment that imposes substantial constraints on cellular proliferation (Harris, 2002), and tested whether a SAPS-defined isoacceptor changed its dependency as predicted under this stress.

Among SAPS-classified isoacceptors, tRNA-Ile-TAT showed one of the strongest restrainer-associated signatures (**Figure 4H**). CRISPRi-mediated depletion of tRNA-Ile-TAT in MDA-MB-231 cells impaired proliferation under both normoxic and hypoxic conditions relative to non-targeting controls (**Figure 4I**, *P* = 2 ×10^−4^), confirming that this isoacceptor contributes to cellular fitness. Importantly, however, the relative fitness defect caused by tRNA-Ile-TAT depletion was significantly attenuated under hypoxia compared with normoxia (**Figure 4J**, *P* = 0.02). Thus, the fitness contribution of an individual tRNA isoacceptor is not fixed, but depends on the environmental state in which its function is required. In particular, tRNA isoacceptor dependency exhibits a tRNA-by-environment interaction: environmental change can quantitatively redistribute the functional requirement for the same component of the decoding machinery. For tRNA-Ile-TAT, this redistribution reduced its requirement under hypoxia, matching the direction expected for a plasticity restrainer. This condition-dependent effect was not observed following perturbation of a cognate tRNA isoacceptor (**Figure 4K**), further indicating that the altered dependency was not a general property of tRNAs decoding the same amino acid, environmental modulation of dependency can distinguish even closely related components of the decoding system. Although hypoxia represents only one component of the complex physiological tumor environment, reproducing the predicted direction of dependency remodeling under a defined stress provides direct experimental support for the central principle emerging from our matched screens: tRNA requirement reflects an interaction between the identity of the decoding component and the physiological environment in which it operates.

### 2.8. Individual tumor-associated stresses induce structured remodeling of tRNA isoacceptor expression

While our functional mapping demonstrates that tRNA dependencies are profoundly reprogrammed by the tumor microenvironment, CRISPRi-based fitness scores report functional requirement rather than tRNA abundance. We therefore asked whether tumor-relevant environmental stresses can also directly remodel the mature tRNA repertoire, and whether this response is shared or context dependent across cellular backgrounds.

To experimentally determine whether tumor microenvironmental stress can directly remodel the mature tRNA repertoire, we exposed *in vitro*–cultured cells to hypoxia. Using MDA-MB-231 and SW480 cells to represent distinct tissue contexts (breast and colon, respectively), we performed mature tRNA sequencing following hypoxic exposure. Principal component analysis of tRNA isoacceptor expression revealed a clear separation between hypoxic and standard conditions (**Figure 5A**; PC2, 29.6% variance explained), with similar response trajectories in both cell lines. Variance decomposition further identified a substantial contribution of hypoxic condition beyond cell identity (**Figure 5B**), demonstrating that individual tumor microenvironmental stress can induce structured remodeling of the mature tRNA repertoire.

**Figure 5.**
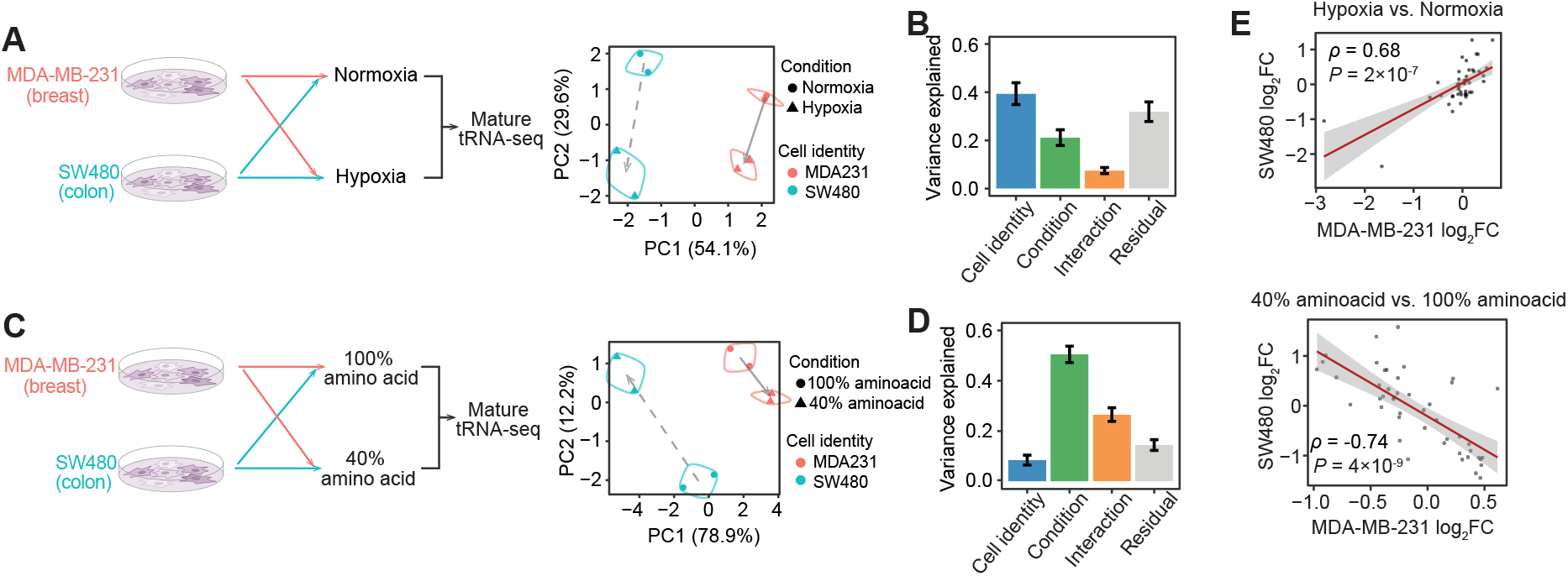
Individual tumor-associated stresses reshape tRNA isoacceptor expression. **(A)** Principal component analysis (PCA) of tRNA isoacceptor expression under standard and hypoxic conditions in MDA-MB-231 and SW480 cells (two biological replicates per condition). **(B)** Fraction of variance in tRNA isoacceptor expression explained by cell identity, hypoxia stress condition, and their interaction. **(C)** Principal component analysis (PCA) of tRNA isoacceptor expression under 100% and 40% amino acid conditions in MDA-MB-231 and SW480 cells (two biological replicates per condition). **(D)** Fraction of variance in tRNA expression explained by cell identity, amino-acid starvation condition, and their interaction. **(E)** Scatter plots comparing changes in tRNA expression between MDA-MB-231 and SW480 cells under hypoxia (upper) and amino acid starvation (lower).

We next asked whether this reprogramming is specific to hypoxia or represents a broader feature of tumorassociated environmental stress. Amino acid limitation represents another major constraint of the tumor microenvironment (Pavlova and Thompson, 2016). Consistent with this, metabolic flux inference from TCGA RNA-seq data revealed widespread signatures of amino acid starvation across tumor types (**Figure S4C**) (Huang et al., 2023; Cancer Genome Atlas Research Network et al., 2013). To model this condition, we cultured MDA-MB-231 and SW480 cells under amino acid–replete (100%) and starved (40%) conditions and profiled mature tRNA abundance (**Figure 5C**). Principal component analysis again revealed coordinated shifts in tRNA expression under amino acid starvation. However, in contrast to hypoxia, these trajectories exhibited cell-type–dependent orientations—remaining parallel in structure but diverging in direction—indicating that tRNA repertoire reprogramming under stress is shaped by both condition and a condition-by-identity interaction (**Figure 5C**). Variance decomposition likewise identified contributions from both environmental condition and its interaction with cell identity (**Figure 5D**).

To directly quantify these distinct response architectures, we compared isoacceptor-level stress responses between the two cell lines in this two stress conditions. Hypoxia-induced changes were strongly concordant between MDA-MB-231 and SW480 cells (Spearman ρ = 0.69, *P* = 2 ×10^−7^; cosine similarity = 0.674; **Fig. 5E**), indicating a broadly shared organization of the hypoxic response. In contrast, responses to amino acid limitation were strongly anti-correlated (Spearman ρ = −0.74, *P* = 4 ×10^−9^; cosine similarity = −0.672; **Fig. 5E**), indicating that the individual component of tumor microenvironment stress can interact with cellular context to reorganize the tRNA repertoire expression. These results suggest that the mature tRNA repertoire is environmentally responsive, but that acute individual stresses can cause different responses interacted with cellular context.

We next asked whether the functional partitioning of tRNA isoacceptors was also reflected in their basal expression across diverse cellular contexts, beyond their responses to individual tumor microenvironmental stress. We therefore analyzed the basal tRNA repertoire across 23 cell lines spanning seven tissues, as characterized in our previous work (Chen et al., 2025). We reasoned that if plasticity restrainers are associated with more stable functional requirements across cellular contexts, their expression might likewise exhibit reduced variation. Indeed, while mean expression levels were comparable across isoacceptors, SAPS scores were significantly negatively correlated with expression heterogeneity (**Figure S8A**). Plasticity restrainers exhibited markedly lower expression heterogeneity (**Figure S8B**), indicating that their expression is comparatively constrained across cellular contexts. Thus, the functional organization identified by SAPS is also reflected in basal tRNA abundance, with restrainer-associated isoacceptors exhibiting greater expression stability across diverse cellular backgrounds. This constrained heterogeneity suggests that plasticity restrainer tRNAs are tightly regulated to support baseline cellular survival and proliferation.

**Figure S8.**
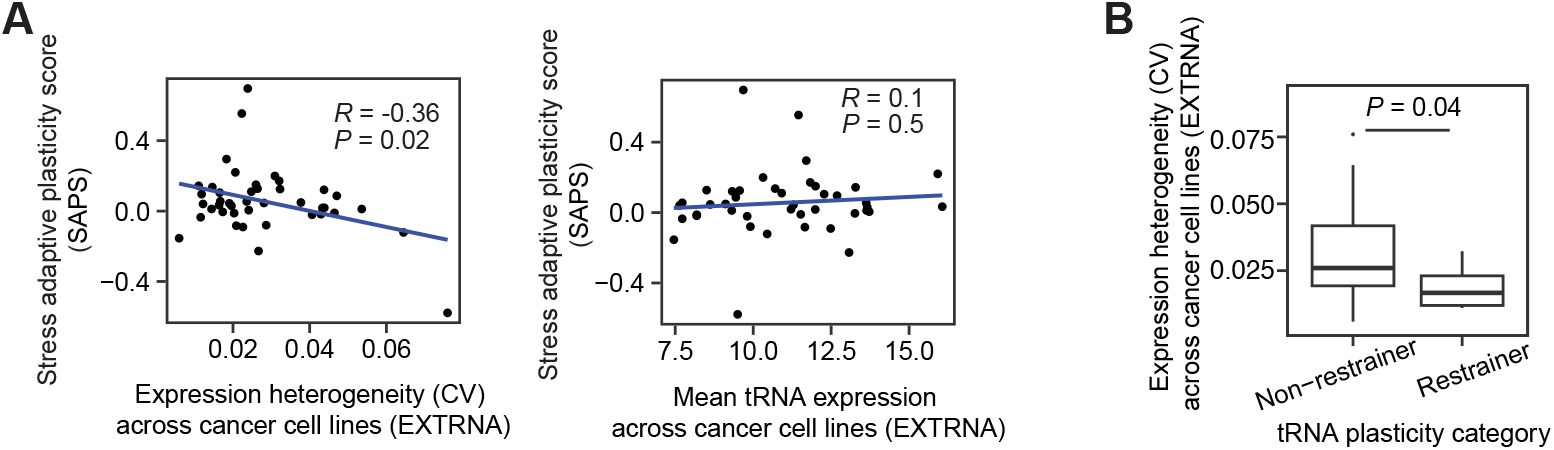
Relationship between stress adaptive plasticity score and basal expression in standard culture in the EXTRNA dataset. **(A)** Scatter plots showing the association between the stress adaptive plasticity score and basal expression heterogeneity (CV) across cells (left) and mean expression levels (right) in the EXTRNA dataset. Pearson correlation coefficients (*R*) and corresponding *P*-values are shown. **(B)** Box plot showing the basal expression heterogeneity (CV) across cells for plasticity restrainer tRNAs compared with other tRNAs. *P*-value was calculated using Mann–Whitney *U* test.

### 2.9. Proliferation and tumor microenvironment–associated stress programs exhibit distinct translational dynamics and codon usage in human tumors

The organization of tRNA isoacceptor dependencies across physiological environments suggested that changes in tRNA requirement may accompany corresponding changes in translational demand. Because individual isoacceptors decode distinct synonymous codons, we asked whether gene programs associated with different cellular states exhibit characteristic codon usage patterns. Analysis of proliferation-, differentiation-, and tumor stress-associated gene sets revealed distinct codon usage patterns, with stress-associated programs showing a particularly different codon composition from proliferation and differentiation programs (**Figure S9**). Thus, functionally distinct cellular programs differ systematically in their codon composition, providing a potential sequence-level basis through which changes in tRNA requirement could be coupled to the translation of state-associated transcriptomes.

We next asked whether these distinct codon demands are accompanied by corresponding differences in translational regulation in human tumors. We leveraged matched CPTAC transcriptome–proteome data (Li et al., 2023; Liao et al., 2023) across four cancer types (BRCA, COAD, LUAD, and PAAD) represented in our *in vivo* models. We estimated per-gene translational output using a multimodal variational framework that accounts for modality-specific variation (**Figure 6A**) (Ashuach et al., 2023). To characterize tumor-associated physiological state, we integrated the translational activities of seven microenvironmental response programs, including hypoxia, inflammatory NF-κB signaling, interferon response, mTOR signaling, oxidative stress, amino acid starvation, and the unfolded protein response. We then separated their shared variation from cancer-type-associated effects and summarized the common component as a Stress Program Index (SPI) (**Figures 6A** and **S10**), representing a major shared axis of tumor physiological variation. Along this coordinate, proliferation and cell-cycle programs exhibited relatively consistent monotonic relationships with translational output, whereas tumor stress-associated programs displayed more dynamic relationships, including sigmoid and pulse-like patterns (**Figure 6B**). Thus, proliferation-associated and tumor microenvironment stress-associated programs differ not only in their codon composition but also in how their translational output varies across physiological states in human tumors.

**Figure 6.**
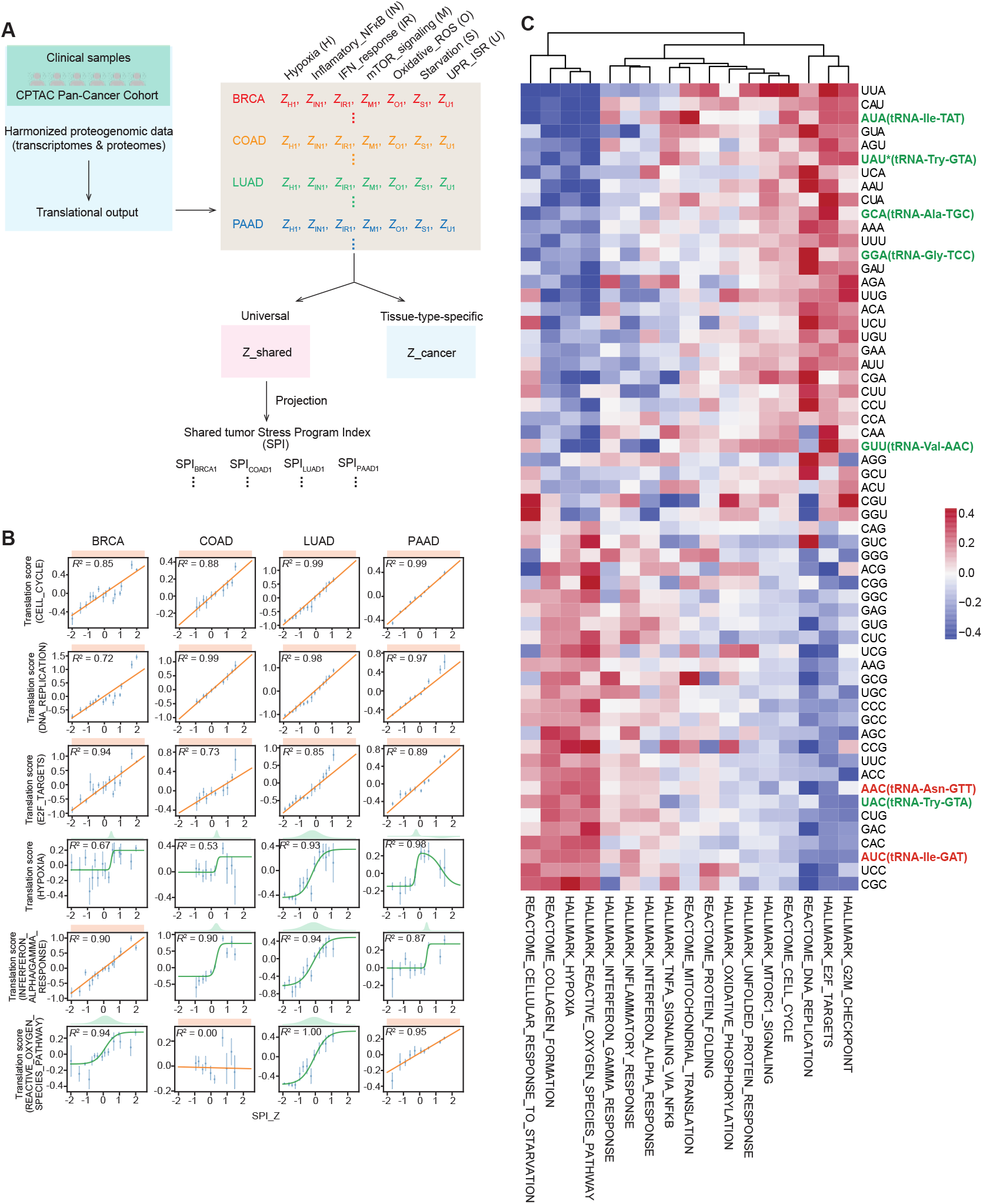
Translational dynamics and codon-usage characteristics of proliferation- and tumor stress-associated gene programs in human tumors. **(A)** Schematic illustrating the unified Stress Program Index (SPI), calculated from CPTAC human clinical samples to represent tumor microenvironment stress. **(B)** Model fitting (constant, sigmoid, and pulse) of translation scores for representative modules (CELL_CYCLE, DNA_REPLICATION, E2F_TARGETS, HYPOXIA, INTERFERON_ALPHA/GAMMA_RESPONSE, REACTIVE_OXYGEN_SPECIES_PATHWAY) along the standardized SPI axis (SPI_Z). **(C)** Heatmap showing codon usage preferences of stress-associated and cell proliferation–associated modules. Genes from **(A)** were used for codon usage analysis, with overlapping genes across terms removed.

**Figure S9.**
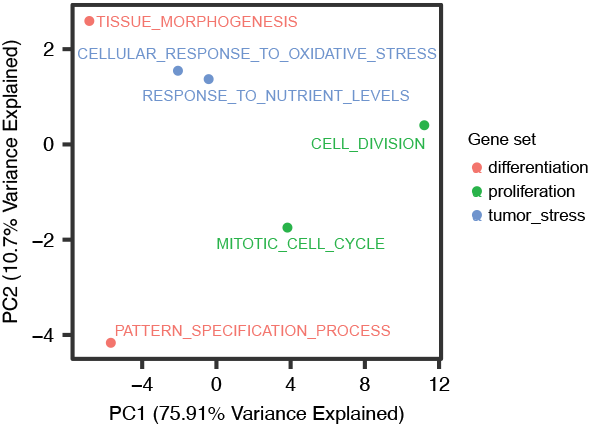
Codon usage preferences associated with differentiation, proliferation, and stress response programs. Principal component analysis (PCA) showing codon usage preferences among genes associated with differentiation, proliferation, and stress response programs. Genes appearing in different GO term were excluded to ensure set-specific comparisons.

**Figure S10.**
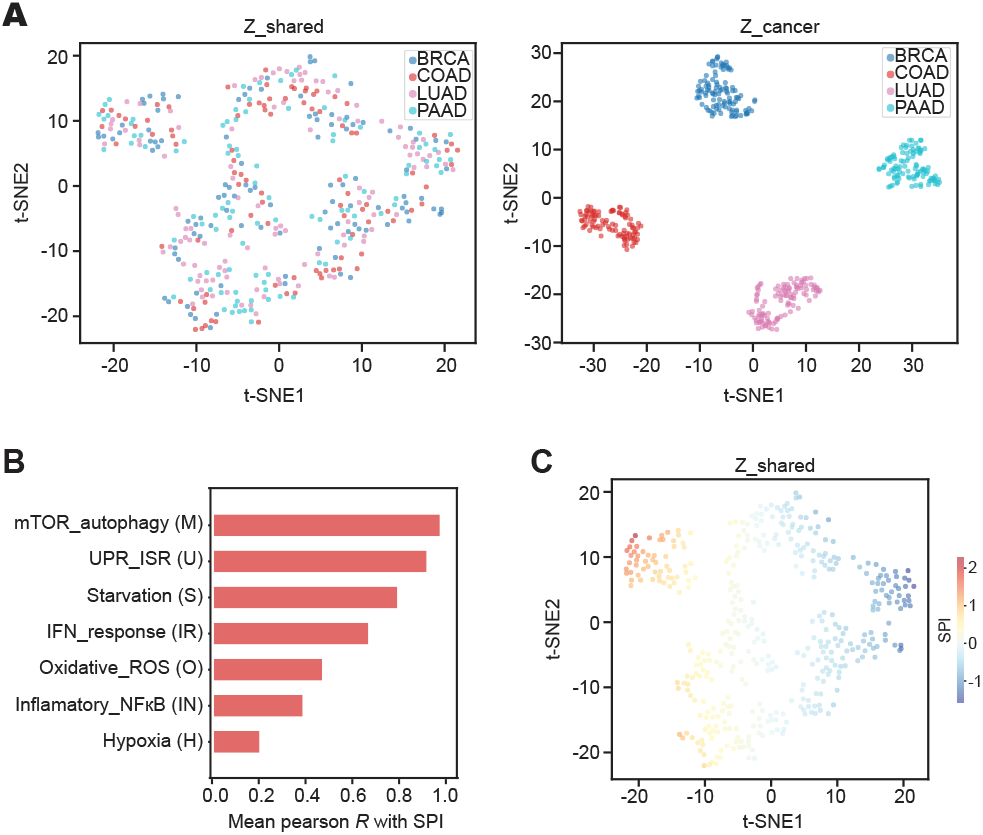
Stress conditions across CPTAC. **(A)** t-SNE visualization of the conditional VAE model, showing separation of the shared latent space (*z*_shared_, left) and the cancer-specific latent space (*z*_cancer_, right) based on module scores of seven stress-associated terms. Points are colored by cancer type (BRCA, COAD, LUAD, and PAAD). **(B)** Bar plot showing the mean correlation coefficients between the calculated SPI and the module scores of each stress-associated term. **(C)** t-SNE visualization showing the SPI gradient in *z*_shared_, with points colored by SPI values.

Furthermore, we asked whether these program-specific codon demands correspond to the tRNA dependency plasticity identified in our functional screens. Notably, codons decoded by plasticity restrainer tRNAs were preferentially represented in proliferation-associated genes, whereas codons decoded by plasticity driver tRNAs were preferentially represented in tumor stress-associated programs (**Figure 6C**). Thus, the functional partitioning of tRNA isoacceptors identified from environmental dependency remodeling is mirrored in the codon composition of distinct translational programs in human tumors. These findings connect the tRNA dependency architecture identified by perturbation to an orthogonal layer of information encoded in transcript sequence, suggesting that environmentally plastic tRNA requirements are coupled to cellular state through differences in codon demand.

### 2.10. Perturbation of the plasticity driver tRNA-Ile-GAT alters translation of tumor stress-associated programs in a codon-dependent manner

Having identified proliferation- and tumor stress-associated programs that clinically exhibited distinct codon usage and translational behavior in human tumors (**Figure 6**), we next asked whether perturbing an individual tRNA isoacceptor identified through our plasticity analysis could directly alter the translational efficiency of the corresponding programs. We reanalyzed matched RNA-seq and ribosome-profiling data from MDA-MB-231 cells in a published study (Earnest-Noble et al., 2022) targeting tRNA-Ile-GAT, which emerged in our analysis as a plasticity driver. tRNA-Ile-GAT depletion caused significant decreases in the translational efficiency of genes enriched for AUC codons decoded by tRNA-Ile-GAT (**Figure S11A**). This indicates that transcripts with greater usage of the decoded codon were specifically affected at the translational level by loss of this isoacceptor.

**Figure S11.**
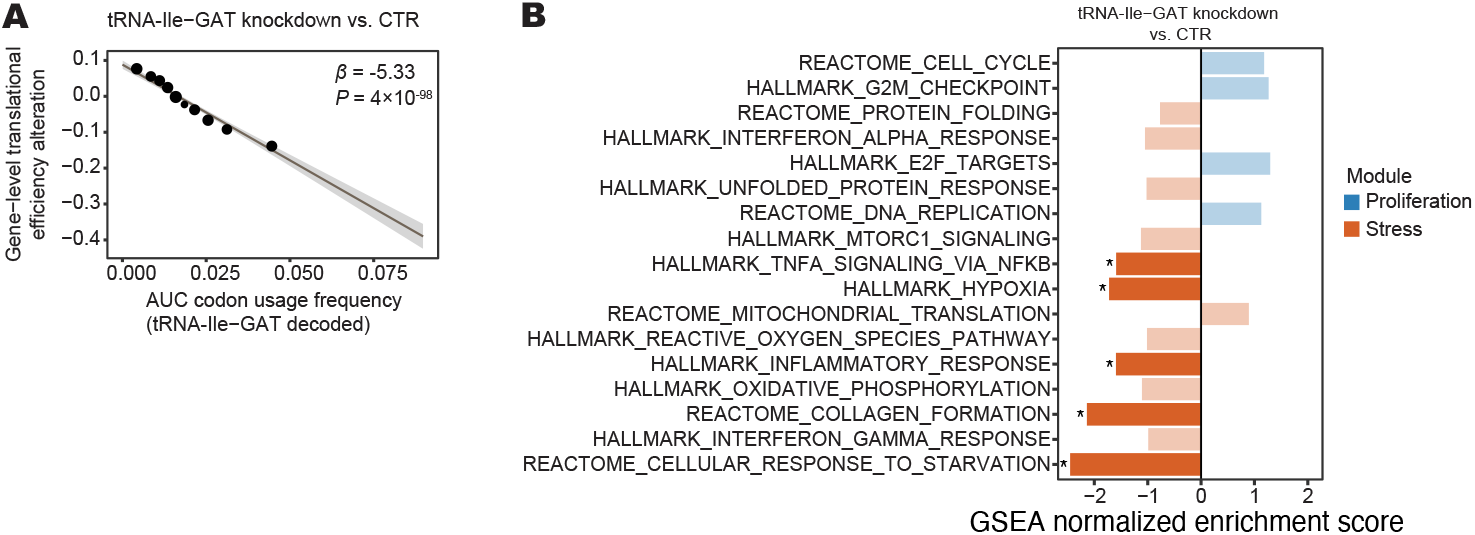
tRNA-Ile-GAT depletion caused codon-dependent translational changes in tumor stress-associated programs. **(A)** Codon usage dependence of translational efficiency alterations following tRNA-Ile-GAT depletion. Differential translational efficiency between tRNA-Ile-GAT knockdown and control cells was estimated from matched RNA-seq and ribosome-profiling data. Points show genes grouped across the codon-usage distribution; the line and shaded region indicate the precision-weighted linear regression and 95% confidence interval, respectively. The regression coefficient (β) and *P* value are shown. **(B)** Pathway-level translational responses following tRNA-Ile-GAT depletion. Bars show normalized enrichment scores, with positive and negative values indicating enrichment toward increased and decreased translational efficiency, respectively. Asterisks indicate pathways with Benjamini–Hochberg-adjusted *P*<0.05

Consistent with our observations in human clinical data, we found that tRNA-Ile-GAT depletion preferentially decreased the translational efficiency of multiple tumor stress-associated pathways, including hypoxia associated and starvation response associated (**Figure S11B**), which were enriched for AUC codon usage (**Figure 6C**). Together with our functional mapping across physiological environments, these findings support a model in which the redundant tRNA repertoire constitutes a structured and environmentally plastic layer of the decoding machinery, coupling changes in tRNA requirement to the translational demands associated with tumor physiological state.

## 3. Discussion

The genetic code is invariant, but the tRNA repertoire that serves as its cellular decoding machinery is not. Redundancy is a defining feature of the tRNA repertoire and has long been associated with translational robustness, yet its functional consequences remain poorly understood, including whether this multiplicity also supports plasticity across physiological environments. Here, systematic perturbation of the cytoplasmic tRNA repertoire across matched *in vivo* tumor growth and *in vitro* cell culture reveals that tRNAs that appear redundant are not functionally equivalent. Instead, tRNA requirements during physiological tumor growth are organized across amino-acid, isoacceptor, and individual isodecoder levels, while this organization exhibits substantial environmental sensitivity that distinguishes tRNAs from the enzymes supporting their maturation and charging. These findings suggest that multiplicity within the tRNA repertoire provides more than essential decoding capacity: it creates a distributed functional architecture in which decoding requirements can be reorganized across physiological environments. Our results therefore suggest that the tRNA repertoire may reconcile two properties of the translational system–robustness of its underlying organization and plasticity of its functional requirements.

A striking feature of this plasticity is that context specificity exists within a highly shared response to physiological environment. During *in vivo* tumor growth, tRNA dependencies exhibited clear tissue organization at isodecoder resolution, whereas this organization was attenuated at the isoacceptor level, where dependencies were instead associated with codon demand. Notably, dependency changes between cell culture and tumor growth were overwhelmingly aligned along a common direction across diverse cancer cells, while the manifestation of this shared response differed across tRNAs and cellular contexts. Separating stable cellular identity from environment-associated dependency states further revealed a shared axis of isoacceptor remodeling and distinguished tRNAs associated with opposing directions of environmental plasticity: plasticity drivers and restrainers. These findings suggest that a common physiological challenge can be accommodated through differential redistribution of requirements among functionally non-equivalent components, allowing substantial remodeling without loss of higher-order organization. Together with the relationship between isoacceptor dependency and codon demand, this architecture positions the tRNA repertoire as a functional interface between decoding requirements and physiological state. In this view, tRNA dependency is not a fixed property of individual molecules, but part of a context-dependent decoding system.

This perspective has broader implications for how coding information is translated into cellular output. At the isoacceptor level, tRNA dependency was associated with codon demand during tumor growth, but this relationship was reorganized in cell culture and further conditioned by the broader demand for the corresponding amino acid. In human tumors, proliferation- and tumor stress-associated programs exhibited distinct codon usage and translational behavior, while independent tRNA perturbation and ribosome-profiling data linked cognate codon usage to changes in translational efficiency and stress-associated pathways. Together, these observations suggest that tRNA dependency plasticity is coupled to the changing translational demands imposed by cellular and physiological state, providing a potential route through which environmental remodeling of decoding requirements can influence translational output. This organization is consistent with previous observations linking synonymous codon usage to distinct biological states and programs (Gingold et al., 2014; Earnest-Noble et al., 2022; Xu et al., 2013). Coding sequence specifies the information presented to the translational machinery, but may not alone determine its functional consequences: the state of the cellular machinery available to decode that sequence also matters. Environmental remodeling of tRNA requirements may thus constitute a functional layer through which the same genetic code is interpreted under different physiological demands.

Our results further reveal an important distinction between environmental organization at the isoacceptor and isodecoder levels. Codon demand provides a natural organizing principle for isoacceptors, but cannot by itself explain the substantial isodecoder-specific structure observed *in vivo*. The enrichment of isodecoder-specific dependencies during tumor growth suggests that elements beyond the anticodon—including body sequence, modification potential, and stability—may contribute to context-dependent physiological requirements. This possibility is supported by mutational studies showing that single-nucleotide changes within tRNA structural elements can have profound functional consequences, with extensive epistatic interactions that are themselves modulated by environmental conditions (Li et al., 2016; Li and Zhang, 2018). Furthermore, the tRNA body serves as a substrate for tissue-specific modification landscapes (Hernandez-Alias et al., 2023; Behrens et al., 2021) and fragmentation pathways that expand the functional repertoire of Pol III-transcribed tRNAs (Huh et al., 2021; Yue et al., 2021; Su et al., 2020; Chen et al., 2021; Goodarzi et al., 2015; Linder et al., 2025; Hughes et al., 2023). Together, these observations support a model in which the tRNA body provides an information-rich layer beyond anticodon identity through which physiological context may shape tRNA function, such that even tRNAs with closely related decoding functions are not functionally interchangeable. Determining how transcription, sequence, modification, charging, stability, and other molecular properties establish this specialization will be important for understanding both the decoding and additional regulatory functions of individual tRNAs across physiological contexts.

Beyond defining these biological principles, our atlas provides a valuable resource for connecting tRNA requirements to the rapidly expanding landscape of context-resolved functional and molecular data. Emerging dependency maps across three-dimensional cultures, patient-derived organoids, and *in vivo* models increasingly demonstrate that physiological context can reshape cellular requirements, extending functional genomics beyond dependencies measured in conventional cell culture (Herranz-Ors et al., 2026; ElHarouni et al., 2026;Neiswender et al., 2026). In parallel, recent findings in spatial profiling, ribosome profiling, and proteomics are revealing substantial context-dependent variation between transcriptional state, translational activity, and protein output across tissues, development, and disease (Yue et al., 2026; Tretyachenko et al., 2026; Tsour et al., 2026). Together, these advances highlight translational regulation as a key context-dependent layer in both gene expression and protein function, through variation in translational efficiency and in the fidelity with which coding information is decoded across physiological states. Understanding the machinery that establishes these context-dependent translational states may therefore be important for explaining how cellular identities and states are established and remodeled across development and disease. Our study connects these emerging directions by adding a largely unexplored functional dimension: the state of the tRNA decoding machinery itself. Our atlas can be integrated with gene-dependency maps to identify pathways associated with particular tRNA requirements; with transcriptomic, translational, proteomic and spatial datasets to investigate where these requirements emerge across cellular states and tissue environments; and with codon-usage, translational, and proteomic measurements to determine how changes in decoding requirements relate to translational efficiency, fidelity, and protein output. Such integration could help connect information across transcriptome, decoding state, translation, and proteome, and reveal where physiological context enters this relationship, and how alterations in this organization contribute to cellular state changes across development and disease. Therefore, our atlas provides not only a reference for tRNA function in cancer, but a foundation for investigating how context-dependent decoding contributes to cellular state transitions across development and disease.

More broadly, our findings emphasize a distinction between molecular state and functional requirement that may extend beyond tRNAs. Molecular abundance alone cannot establish whether an individual component is dispensable, buffered by related molecules, or selectively required in a particular environment. The extensive reorganization of tRNA dependencies between cell culture and tumor growth further demonstrates that essentiality measured under a single condition need not represent a fixed property of a gene or noncoding element. This distinction may be particularly important for highly redundant and comparatively unexplored noncoding systems, in which apparent redundancy can conceal functional specialization until the appropriate physiological context is encountered. As functional genomics increasingly moves into physiologically resolved systems, systematic perturbation of noncoding elements across cellular states and environments may uncover layers of biological organization that cannot be inferred from sequence, molecular abundance, or conventional protein-coding dependency maps alone.

Moving forward, our observation of context-selective tRNA dependencies raises the possibility of therapeutic vulnerabilities. Specific tRNAs—or the regulatory factors responsible for establishing their selective dependencies—may provide opportunities to perturb translational requirements that become particularly important in defined physiological states. In particular, plasticity-associated tRNAs provide candidates for testing whether disruption of environmentally adaptive decoding requirements can constrain tumor adaptation. Establishing such therapeutic potential, however, will require determining the selectivity of these dependencies across normal tissues and whether they can be targeted without compromising essential translation.

## 4. Limitations

Several aspects of the functional architecture identified here remain unresolved. Our *in vivo* experiments model tumor growth across a diverse panel of cancer cell lines but do not capture the full range of environments encountered by tumors in patients. Xenograft growth provides a physiologically complex *in vivo* context but does not fully reproduce the tissue architecture, immune environment, or evolutionary history of native human tumors. Moreover, the relative contributions of nutrient availability, oxygen tension, stromal interactions, immune signals, and other physiological factors to the observed dependency remodeling remain to be resolved. Our defined stress experiments demonstrate that individual environmental pressures can remodel tRNA dependency and expression, while also showing that distinct stresses need not produce equivalent tRNA responses. Expanding functional tRNA mapping across additional physiological states, normal tissues, and defined environmental perturbations will therefore be important for determining how broadly the principles identified here extend beyond the tumor models examined.

Our analyses further identify a shared axis that organizes tRNA dependency remodeling between *in vitro* and *in vivo* environments, but they do not establish this axis as a temporal trajectory of tumor adaptation. Likewise, the association of individual tRNAs with opposing directions along this axis provides a framework for identifying plasticity drivers and restrainers but does not by itself establish the molecular mechanisms through which these tRNAs influence adaptive cellular states. Longitudinal perturbation studies and direct measurements of downstream cellular states will be required to determine how movement along this axis relates to adaptation over time and whether individual tRNAs causally promote or constrain progression along this adaptive state space. The molecular determinants underlying these differential responses also remain unresolved, particularly at isodecoder resolution. Direct measurements of tRNA transcription, modification, aminoacylation, turnover, and molecular interactions across defined physiological contexts will be needed to determine how these regulatory layers establish the context-dependent dependencies identified here.

## 5. Methods

All animal experiments were conducted in accordance with relevant ethical regulations and were approved by the UCSF Institutional Animal Care and Use Committee (IACUC, protocol AN179718).

### 5.1. Cell culture

All cell lines were maintained at 37°C in a humidified incubator with 5% CO_2_. HEK293T (ATCC CRL-3216), MDA-MB-231 (ATCC HTB-26), A375 (ATCC CRL-1619), MIA PaCa-2 (ATCC CRL-1420), A2058 (ATCC CRL-3601), MDA-MB-453 (ATCC HTB-131) and HCC1806 (ATCC CRL-2335) cells were cultured in high-glucose DMEM supplemented with 1 mM sodium pyruvate. PC-3 (ATCC CRL-1435) cells were cultured in F-12 medium. HCT116 (ATCC CCL-247) and SW480 (ATCC CCL-228) cells were cultured in McCoy’s 5A medium. C4-2B (ATCC CRL-3315), H23 (ATCC CRL-5800), K562 (ATCC CCL-243), COLO320 (ATCC CCL-220), H1299 (ATCC CRL-5803), H358 (ATCC CRL-5807), and HCC44 (DSMZ ACC-534) cells were cultured in RPMI-1640 medium supplemented with 25 mM HEPES. All media were supplemented with 2 mM L-glutamine, 10% fetal bovine serum (FBS), 100 U/mL penicillin, 100 μg/mL streptomycin, and 1 μg/mL amphotericin B (Gibco). All cell lines were routinely tested for mycoplasma contamination using a PCR-based assay and confirmed negative.

### 5.2. Generation and validation of CRISPRi cell lines

Lentiviral particles carrying dCas9-ZIM3-KRAB (Alerasool et al., 2020) were produced by co-transfecting the dCas9-ZIM3-KRAB construct with pCMV-dR8.91 and pMD2.D packaging plasmids into HEK293T cells using TransIT-Lenti (Mirus) following the manufacturer’s protocol. Virus-containing supernatants were collected 48 h post-transfection, filtered through a 0.45 μm filter, and used to transduce target cell lines in the presence of 8 μg/mL polybrene (Millipore). Transduced cells were enriched by fluorescence-activated cell sorting (FACS) for GFP-positive populations twice to establish stable CRISPRi cell lines. All cell identities were confirmed with STR analyses, and tested for mycoplasma contamination using a PCR-based assay and confirmed negative.

To assess the knockdown efficiency of generated CRISPRi cell lines, we introduced sgRNAs targeting ST3GAL4, ERK1, and SEL1L individually (Horlbeck et al., 2016). sgRNA sequences were cloned into the pCRISPRia-v2 vector (Addgene 84832) via BstXI–BlpI sites. After lentiviral transduction, cells were selected with 1.5–3 μg/mLpuromycin (Gibco). Knockdown efficiency was quantified by RT-qPCR, comparing expression levels to those in control cells transduced with a non-targeting guide (gCTR).

### 5.3. Lentiviral guide RNA library generation

Guide RNA sequences targeting actively transcribed human tRNAs were designed using FlashFry (McKenna and Shendure, 2018). The library also included sgRNAs targeting aminoacyl-tRNA synthetases (aaRSs) and tRNA-modifying enzymes, and 100 non-targeting control guides. The oligo pool was synthesized by Twist Bioscience and PCR-amplified using primers SC262 and SC263 with Phusion High-Fidelity DNA Polymerase (NEB). Amplified products were purified using 2% agarose gel extraction and cloned into the pBA904_gba405 vector via BstXI–BlpI sites using NEBuilder HiFi DNA Assembly Master Mix (NEB). Ligation products were ethanol-precipitated, electroporated into Meta-x, and plated on ampicillin selection plates. The resulting plasmid pool was verified by amplicon sequencing.

For lentivirus production, the plasmid pool was co-transfected with pCMV-dR8.91 and pMD2.D into HEK293T cells using TransIT-Lenti (Mirus). Viral supernatants were harvested 48 h post-transfection, filtered (0.45 μm), aliquoted, and stored at −80°C.

### 5.4. tRNA CRISPRi in vitro screen

Prior to large-scale screening, viral titration experiments were performed for each cell line by transducing cells with varying viral volumes. Flow cytometry was used to determine the optimal volume that achieved a transduction efficiency of approximately 10–20%, which was then applied for screening experiments. Transduced cells were selected with puromycin (1.5–3 μg/mL) until all untransduced control cells (no sgRNA lentivirus) were eliminated. Following selection, cells were cultured in puromycin-free medium for 1–2 days before initiating the screen. At *T*_0_, cells were divided into three groups:

1. 2 × 10^6^ cells were collected by centrifugation and frozen for genomic DNA extraction (*T*_0_ library).
2. 7 × 10^6^ cells were prepared for in vivo screening.
3. 2 × 10^6^ cells were seeded in 10 cm plates to initiate in vitro screening.

Cells were passaged every other day, and cell pellets were collected at multiple time points (e.g., Days 8, 10, and 12; or adjusted based on growth rate). Genomic DNA was extracted using the Quick-DNA Miniprep Kit (Zymo Research).

### 5.5. tRNA CRISPRi in vivo screen

For each cell line, 6 ×10^6^ cells were injected for in vivo screening in NSG mice. Cultured cells were harvested by trypsinization and centrifugation, washed twice with sterile 1 × PBS to remove residual medium, and resuspended for injection. For breast cancer cell lines, each mouse received two injections into the mammary fat pads, one on each side, with 1.5 × 10^6^ cells per site suspended in 100 μL of a 1:1 mixture of 1 × PBS and Matrigel (Corning). For other cancer cell lines, injections were performed subcutaneously into each flank, with 0.8 ×10^6^ cells per site in 100 μL of the same 1:1 PBS/Matrigel mixture. Mice were euthanized in accordance with approved IACUC protocols once tumors reached a volume of approximately 1,000 *mm*^3^. Tumors were aseptically excised, weighed, and immediately processed for downstream molecular analyses. Genomic DNA was isolated from freshly dissected tumor tissue using the Quick-DNA Midiprep Plus Kit (Zymo Research), following the manufacturer’s instructions.

### 5.5. UMI-linked guide RNA library construction and sequencing

Extracted genomic DNA was digested with BstXI and ligated to a UMI-containing linker using Klenow (exo-) and 50% PEG8000. Ligated fragments were purified with DNA clean & concentration-25 column (Zymo Research) and amplified with Phusion polymerase, followed by gel purification using a 4% agarose gel. The amplified UMI-linked guide libraries were sequenced in SE100 mode on the Illumina NovaSeqX instrument at UCSF Center for Advanced Technologies.

### 5.6. Calculation of guide RNA growth rate and normalization

Sequencing reads were first processed to extract unique molecular identifiers (UMIs) using umi_tools (`v1.1.5; –bc-pattern=NNNNNNXXXXXXX`), followed by adapter trimming with cutadapt (`v4.9; -m 19 -M 21 -q 15 -a ^CTTGTTG…GTTTAAGAGCTAAGCTG`). The trimmed reads were then aligned to the pooled guide RNA reference FASTA file using bowtie2 (v2.5.4; –sensitive –end-to-end -N 1). PCR duplicates were removed based on UMI information, and the remaining unique sgRNA reads were counted. For sgRNAs that can be mapped to multiple tRNA isodecoders, isoacceptors, or amino acid families, the guide sequences were annotated with all possible family identities in a reference file rather than being assigned to a single one.

For each cell line, DESeq2-style size factors were first computed to normalize sequencing depth. Specifically, for each guide RNA, the geometric mean of counts across samples was calculated, and the median ratio of observed counts to this geometric mean was used as the sample-specific size factor after removing zero and non-finite values. Normalized counts were then obtained by dividing raw counts by these size factors.

To ensure the reliability of downstream analyses, stringent quality control (QC) was performed at both the guide and timepoint levels. For each guide in each cell line, the following metrics were computed: (1) presence of a baseline (*T*_0_) measurement and its read count, (2) number of available and sufficiently abundant timepoints (≥12 raw counts), (3) extreme fold changes across timepoints (flagged if > 50×, and (4) variability of log-transformed normalized counts over time, measured as the standard deviation of residuals from a linear regression (flagged if > 1.5). Guides were classified as problematic guides if they met any of the following criteria: missing or low-count *T*_0_, fewer than two usable later timepoints, excessive fold change, or high variability. These guides were excluded from subsequent analyses.

For each remaining guide, timepoint-level QC was further applied to detect and remove outlier observations. For each guide trajectory, a linear model of log-normalized counts versus time was fitted (excluding *T*_0_ and low-count timepoints), and individual timepoints with residuals exceeding an absolute threshold (> 1.0) were flagged as outliers and excluded, while *T*_0_ samples were always retained.

Following QC filtering, guide-level growth rates were estimated independently within each cell line. A robust linear regression (rlm) was applied to log-transformed normalized counts relative to *T*_0_ using the model:

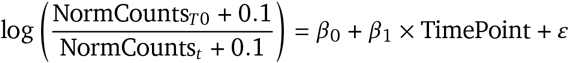

A small pseudocount (0.1) was added to avoid undefined log values. To correct for global effects and facilitate comparability across cell lines, guide-level growth rates were normalized using non-targeting control guides. For each cell line, the median and median absolute deviation (MAD) of non-targeting guide growth rates were computed, and all guide scores were transformed into robust z-scores with with (growth_rate −Median_NT_)/MAD_NT_.

Processing of in vivo CRISPRi screen data followed the same general framework as the in vitro analysis but with simplified time structure and QC criteria. Because each in vivo screen contained only two timepoints (*T*_0_ and *T*_*t*_), timepoint-level outlier detection was not performed. Instead, only guides that passed in vitro QC and were also present in the in vivo dataset were retained for comparison. For each remaining guide, growth rate was calculated directly as the log fold change of normalized counts between *T*_*t*_ and *T*_0_.

To correct for experimental noise, batch effects and cross-cell line variability in guide RNA growth rate estimates, we employed a variational autoencoder on guide-level growth rate results. The model was trained with biologically guided constraints on the distributions of non-targeting guides and essential guides (essential genes defined from DepMap). Non-targeting guides were constrained to follow a normal distribution centered at 0 (*σ* ≈ 0.5), while essential guides were constrained to have a consistent depletion effect (mean *logFC* < −1.25). Additionally, a false-negative rate (FNR) penalty ensured that the empirical overlap between essential and non-targeting distributions remained below 0.05. To preserve learned noise structure while adapting to *in vivo* conditions, the pretrained VAE was fine-tuned for *in vivo* dataset using the same architecture and loss components. Considering we only have negative anchors (essential guides) were available, which introduce asymmetry, we further employed a Variance Stabilization (VST) procedure on denoised guide RNA growth rate to balance the distribution.

To control for off-target effects, we re-annotated all sgRNAs using CRISPOR (hg38, SpCas9/NGG) (Concordet and Haeussler, 2018). We focused on high-confidence predicted off-target sites (≤1 mismatch) and intersected these with protein-coding transcript annotations (GENCODE v44). sgRNAs were excluded if any predicted off-target site overlapped a protein-coding promoter (defined as ± 5 kb around the annotated transcription start site) or protein-coding transcript body outside the intended tRNA locus.

### 5.8. Perturbation score calculation of tRNA molecule and protein coding gene

Normalized guide-level growth rates were next aggregated to the molecule level and gene level to estimate perturbation effects for each targeted molecule. For each cell line, gene-level scores were summarized using the 10% trimmed mean of constituent guides to reduce outlier influence. Minimum guide thresholds were applied: ≥2 guides for tRNA molecules and ≥3 guides for protein-coding genes. Genes below these thresholds were excluded. For analyses at the tRNA isodecoder and isoacceptor levels, only guides that uniquely target a single isodecoder or isoacceptor were retained.

Empirical *P*-values were computed from a null distribution of pseudo-genes generated from non-targeting guides. Specifically, for each cell line, non-targeting guides were randomly grouped into pseudo-genes of guide RNA sizes 2–6 to match the real guide-count distribution. Then, for 10,000 bootstrap iterations, 10% trimmed means were calculated to generate a null distribution of expected scores. Two-tailed empirical *P*-values were calculated based on the obtained null distribution.

### 5.9. Individual tRNA knockdown and quantification by RT–qPCR

Individual tRNA knockdowns were performed as previously described. Non-targeting control guide sequences were obtained from a published study (Replogle et al., 2022). Single-guide plasmids were co-transfected with the packaging plasmids pCMV–dR8.91 and pMD2.D into HEK293T cells using TransIT-Lenti (Mirus) according to the manufacturer’s instructions. Virus-containing supernatants were collected 48 h post-transfection, filtered through a 0.45 μm filter, and used to transduce target cells in the presence of 8 μg/mL polybrene (Millipore). Transduced cells were selected with 2 μg/mL puromycin until all untransduced control cells (no sgRNA lentivirus) were eliminated.

Total RNA was extracted from selected cells using the Quick-RNA kit (Zymo Research). Relative tRNA expression levels were quantified by RT–qPCR. Briefly, total RNA was reverse transcribed to cDNA using SuperScript III Reverse Transcriptase (Thermo Fisher Scientific), followed by amplification with PowerUp SYBR Master Mix (Applied Biosystems) according to the manufacturer’s instructions. 5S rRNA served as the endogenous control. Two biological replicates, each measured in four technical replicates, were performed in RT-qPCR.

### 5.10. Tissue classification with tRNA isodecoder CRISPRi perturbation scores

An input matrix was constructed with samples as rows and tRNA isodecoder perturbation scores as features. For each feature, the median and median absolute deviation (MAD) were computed across samples. To remove low-variance features, only those with MAD values above the 25th percentile of the overall MAD distribution were retained.The remaining features were robustly standardized as 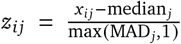 where MAD values equal to zero were replaced with 1 to prevent division by zero.

To capture the dominant structure of tRNA perturbation patterns without label leakage, principal component analysis (PCA) was fitted once on the robustly scaled data using all samples without tissue labels. The top three principal components were retained for downstream analyses. This PCA basis and corresponding scaling parameters were held fixed in all cross-validation and permutation steps. Only tissues with at least three samples were included in the evaluation.

Tissue classification was performed in this fixed PC space using shrinkage linear discriminant analysis (LDA). Model performance was assessed by leave-one-out cross-validation (LOOCV), in which the PCA projection remained fixed and only the LDA classifier was refitted per fold using *n* −1 samples. To evaluate whether observed accuracy exceeded chance, tissue labels were randomly permuted (class sizes preserved) across *N* = 10,000 iterations while keeping the PCA basis fixed. The one-sided empirical *P*-value was computed as the fraction of permutation accuracies greater than or equal to the observed accuracy.

To identify tRNA isodecoders contributing to tissue separation, one-vs-rest LDA contrasts were calculated in the fixed PC space and mapped back to the original robust-scaled feature space using PCA loadings, yielding signed coefficients for each feature and tissue. Feature importance was defined as the absolute coefficient magnitude. Empirical *P*-values for each feature–tissue pair were obtained by repeating the label permutation procedure (*N* = 10,000) with the PCA basis fixed and recomputing the corresponding contrasts. Features were considered significantly associated with tissue separation at *P* ≤ 0.05.

### 5.11. Multi-tissue self-attention with contrastive learning for tissue-specific tRNA isodecoder coordination discovery

Sample metadata (tissue labels) were merged with the tRNA isodecoder perturbation matrix. For model training and validation, features were standardized by a z-score transform fit globally across all samples (StandardScaler).

A multi-tissue self-attention reconstruction model was employed that jointly learns (i) tRNA isodecoder interaction patterns and (ii) a low-dimensional contrastive embedding aligned with tissue identity. Each tRNA is treated as a token with a learnable identity embedding; per-sample scalar values are projected to the hidden space and concatenated with (i) the tRNA embedding and (ii) a learnable *tissue* embedding. The concatenated representation passes through a combining layer (ReLU, dropout), followed by single-head self-attention and an output projection that reconstructs the input profile. Two auxiliary heads are included: a tissue-classification head (cross-entropy loss) on the pooled token representations, and a contrastive-projection head that embeds the flattened attention matrix to support supervised contrastive learning.

Parameters were optimized with Adam to minimize a weighted sum of reconstruction, classification, and contrastive losses:

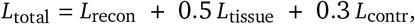

where *L*_recon_ is mean squared error between reconstructed and observed profiles, *L*_tissue_ is cross-entropy for tissue prediction, and *L*_contr_ is a supervised contrastive loss applied to the contrastive projection of the attention matrix, encouraging samples from the same tissue to cluster in embedding space while separating different tissues.

Tissues represented by at least three cell lines were included in LOO validation. For a given tissue, the stability of each tRNA isodecoder pair (*i, j*) was quantified across LOO folds as

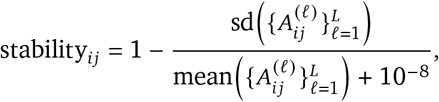

where 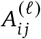 denotes the attention value for pair *i, j* in the tissue-level attention matrix estimated in LOO fold *l*, and *L* is the number of folds. A tissue-specific null distribution was constructed by permuting tissue labels globally across all samples (*N* = 10,000 permutations). For each permutation, the scaler was refit globally, the model retrained, and per-tissue coordination matrices were computed by averaging attention matrices within the permuted tissue. Empirical two-sided *p*-values for each pair *i, j* and tissue were computed as the fraction of null values with absolute magnitude at least as large as the observed coordination. Significant tRNA isodecoder-isodecoder pairs were defined as those with *P* < 0.05 and stability > 0.7.

### 5.12. Chromatin accessibility–matched decoy pair construction using TCGA ATAC-seq

Chromatin-matched decoy pairs were constructed using TCGA pan-cancer ATAC-seq data (log-normalized peak counts) (Corces et al., 2018). BRCA, COAD, and LUAD were used in this analysis, corresponding to the tissues included in our multi-tissue self-attention analyses.

The full set of tRNA isodecoder pairs was used as the reference score table. Human tRNA loci were obtained from the hg38 tRNA annotation BED file, and loci overlapping ENCODE blacklist regions (ENCFF356LFX) were excluded to retain loci with interpretable chromatin accessibility signals.

To summarize chromatin accessibility at each tRNA locus, ATAC-seq peaks overlapping a ± 500 kb genomic window centered on each locus were identified using genomic interval overlaps. For each tRNA locus, the mean signal across overlapping peaks was calculated for each TCGA sample. Tissue-specific chromatin accessibility was then summarized as

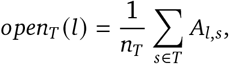

where *A*_*l,s*_ denotes the ATAC-seq signal of locus *l* in sample *s*, and *n*_*T*_ is the number of samples in tissue *T*.

For each significant tRNA isodecoder pair (*i, j*) within a tissue (identified from the multi-tissue self-attention analyses), we constructed a decoy set for partner *j* while preserving the anchor isodecoder *i*. Specifically, all loci belonging to isodecoder *j* were considered. Candidate decoy loci *k* were required to satisfy the following constraints: matched copy class, matched peak-count bin, similar tissue-specific chromatin accessibility,and exclusion of loci belonging to real partners of anchor *i* or to isodecoder *j* itself.

Candidate decoys identified across all loci of *j* were pooled and collapsed to unique decoy isodecoders by retaining the best accessibility match. Up to 100 decoy isodecoders were then sampled for each real pair to construct the chromatin-matched null set.

### 5.13. H3K4me3 analysis of transcriptional activity at tRNA loci

H3K4me3 fold-change-over-control bigWig tracks and replicated narrowPeak calls were obtained from the ENCODE portal (v4) (ENCODE Project Consortium, 2012) for three cell lines representing the tissues analyzed in our multi-tissue self-attention model: MCF7 (breast; ENCFF935BFQ / ENCFF992GMR), A549 (lung; ENCFF343IDC / ENCFF753WFU), and HCT116 (colon; ENCFF964OOU / ENCFF187LLD), all aligned to GRCh38. For each tRNA gene locus, the mean H3K4me3 fold-change signal was extracted from the bigWig track within a ± 500 bp window centered on the locus midpoint. A locus was considered transcriptionally active if the mean fold-change exceeded a threshold of 2.0.

For each significant tRNA isodecoder pair (*i, j*) in a given tissue, all pairwise combinations of genomic loci belonging to isodecoders *i* and *j* were enumerated. A gene-level pair was classified as *both-active* if both constituent loci exceeded the H3K4me3 activity threshold in the matched cell line.

### 5.14. Paired context-aware variational autoencoder (pcVAE) in cell identity-state separation

For each paired sample between in vitro and in vivo perturbations 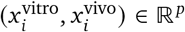, a shared encoder maps input features into two latent representations: a condition-invariant *identity* component and a condition-specific *state* component,

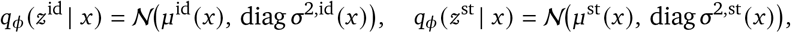

where reparameterization is applied as *z* = μ *σ* ⊙ϵ, ϵ ∼ N (0, *I*) . A decoder *g*_θ_ reconstructs the original input from the concatenated latent vector [*z*^id^ ∥*z*^st^] . The dimensionalities of *z*^id^ and *z*^st^ were guided by principal component analysis (PCA) performed on two transformed matrices: the sum (*x*^vivo^ +*x*^vitro^), representing shared variation across contexts, and the difference (*x*^vivo^−*x*^vitro^), representing context-specific variation. The number of latent dimensions for *z*^id^ and *z*^st^ was chosen as the smallest number of principal components jointly explaining at least 95% of the total variance in each respective space.

The pcVAE model was trained to jointly optimize reconstruction accuracy, identity consistency, state separation, orthogonality between latent subspaces, and KL regularization:

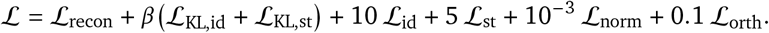

#### Reconstruction

Reconstruction fidelity was measured using the Smooth L1 (Huber) loss for both conditions:

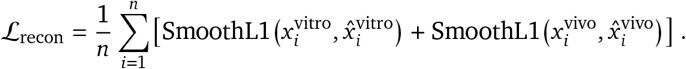

#### Identity alignment

To ensure that identity representations remain consistent across in vitro and in vivo conditions, the distance between paired identity latents was penalized:

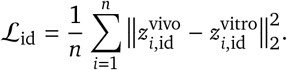

#### State separation

To enforce environmental specificity, state latents were encouraged to diverge between paired conditions using a hinge loss:

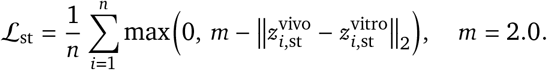

#### Orthogonality constraint

To reduce redundancy between identity and state spaces, the cross-covariance of their centered representations was penalized:

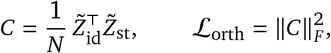

where 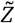 denotes column-centered latent matrices across both conditions.

#### State norm penalty and KL regularization

To prevent trivial inflation of state magnitudes, we included a mild *L*_2_ penalty,

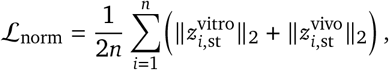

and applied standard Gaussian KL divergence to both posteriors, annealing β linearly to 0.01 during training.

The pcVAE was trained using Adam (learning rate 10^−3^, weight decay 10^−4^) with gradient clipping (global norm = 1.0) and early stopping. The selection of the model was based on a composite validation metric combining reconstruction loss, identity consistency, and state separation. Final weights were checkpointed and restored for downstream analyses.

### 5.15. Disentanglement validation by identity–state swap reconstruction

To assess the quality of disentanglement achieved by the pcVAE model, we evaluated both representation consistency and functional independence between identity and state latent spaces. For each matched in vitro– in vivo pair, the condition-invariant (identity) and condition-specific (state) embeddings were denoted as 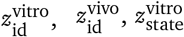, and 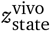, respectively.

Identity preservation was quantified as the average Euclidean distance 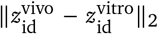, while state separation was computed as the mean norm of 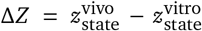 Their ratio provided a scalar measure of disentanglement tightness, with smaller values indicating clearer separation of invariant and adaptive components.

To functionally validate independence between the two subspaces, we performed an identity–state *swap reconstruction* test. For each sample *i*, its true identity embedding 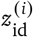 was replaced with that of another sample 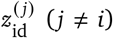, while keeping its state embedding 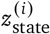 unchanged. The decoder then reconstructed the original input from the swapped latent combination 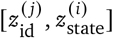. The resulting mean squared error (MSE) was compared with the reconstruction error using the correct pairing, and the swap ratio (MSE_swap_ /MSE_original_) quantified the extent of disentanglement. Larger swap ratios indicated successful orthogonalization of identity and state representations, where mismatched latent compositions degraded reconstruction accuracy.

### 5.16. Variance decomposition of tumor–environment adaptation

Variance components associated with adaptation from in vitro to in vivo were decomposed in two stages: a *system-level* analysis on latent state shifts (Δ*z*) and a *feature-level* analysis on decoder-space changes (Δ*x*). The inputs were standardized with a single scaler fit to the row-wise concatenation of in vitro and in vivo matrices ([*X*^vitro^; *X* ^vivo^]) prior to model fitting and downstream analyses.

For each cell line *i*, identity and state latents were obtained for both conditions, and two derived quantities were computed:

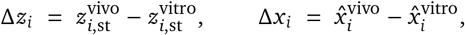

Where 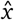 denotes pcVAE reconstructions. Condition-invariant identity was summarized as 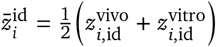.

To capture baseline genetic structure without leaking outcome labels, principal components of identity were computed on *z*-scored 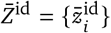. The smallest number of components (*idPCs*) jointly explaining at least 80% of variance was retained. A global “shared-stress” score per cell line was extracted from Δ*Z* = {Δ*z*_*i*_ }as the first principal component (1D projection), then *z*-standardized.

#### System-level variance on Δ*z*

Latent state shifts were analyzed with an ANOVA including (i) tissue effects and (ii) idPC-by-tissue interactions to modulate tissue effects by baseline identity. Tissues with fewer than three samples were collapsed into an Other level for the tissue terms. For each latent dimension *k*,

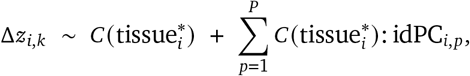

and sums of squares were converted to proportions of variance (*h*^2^) for tissue, gene tissue (idPC × tissue) interaction, and residual components.

#### Feature-level variance on Δ*x*

For each tRNA feature *j*, the decoder-space change was modeled as

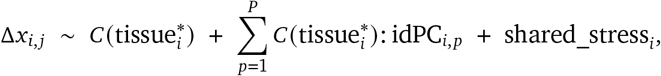

using the same tissue-filtering rule described above. For each feature, proportions of variance (*h*^2^) were obtained for the tissue main effect, the gene × tissue (idPC-by-tissue) interaction, the shared-stress covariate, and the residual. Statistical significance was evaluated by a component-wise permutation test (*N*=10,000) using right-tailed empirical *p*-values (fraction of permuted *h*^2^ exceeding the observed). Components with *p*<0.1 were considered significant.

### 5.17. Calculation of Stress Adaptive Plasticity Score (SAPS) and category classification

For each cell line *i*, the pcVAE provided state latent shifts 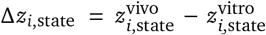. A global stress axis *u* was defined as the first principal component (PC1) of all Δ*zi*,state The sign of *u* was aligned to ensure anonnegative mean projection. Each cell line’s stress severity was then computed as its projection onto this axis and standardized to unit variance:

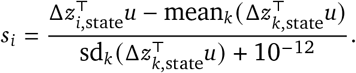

For each tRNA feature *j*, decoder-space perturbation shifts were obtained as 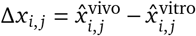. The SAPS was defined as the regression slope between Δ*x*_*i, j*_ and the standardized stress severity:

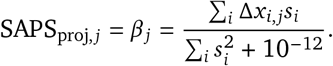

This slope quantifies how strongly each tRNA feature’s perturbation effect scales with the inferred stress level in tumor adaptation.

### 5.18. Proliferation rate quantification

Individual tRNA knockdowns were performed in MDA-MB-231 cells using a dual-guide CRISPRi system as described above. Cells were seeded at a density of 1,000 cells per well in 100 μL of growth medium in 96-well tissue culture plates (Corning, 3695), with eight biological replicates per condition (gCTR, gtRNA-Ile-TAT, gtRNA-Ile-AAT).

Cells were cultured under either standard conditions (21% O_2_) or hypoxic conditions (1% O_2_, 5% CO_2_, balance N_2_). Cell proliferation was quantified using the CellTiter-Glo Luminescent Cell Viability Assay (Promega, G7570) according to the manufacturer’s instructions.

For each oxygen condition, independent replicate plates were prepared to enable time-resolved measurements of growth dynamics. Under standard conditions, luminescence was recorded as Day 1, 2, 4, and 5. Under hypoxia, cells were pre-incubated for 48 hours to ensure complete adaptation to low oxygen before growth measurements were initiated, and luminescence was subsequently recorded as Day 1, 2, and 4.

Background-corrected luminescence signals were used to estimate proliferation rates by fitting an exponential growth model:

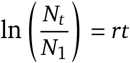

where *N*_*t*_ and *N*_1_ represent luminescence values at Day *t* and Day 1, respectively, and *r* denotes the proliferation rate.

### 5.19. Metabolic flux inference from TCGA RNA-seq data

TCGA RNA-seq expression matrices (Cancer Genome Atlas Research Network et al., 2013) were processed using the METAFlux framework (Huang et al., 2023) to infer metabolic reaction activity and estimate steady-state fluxes under a human blood–like medium. Amino acid–related fluxes were extracted based on nutrient category annotation, where positive values represent predicted metabolite release and negative values represent uptake.

### 5.20. Mature tRNA library sequencing under hypoxia and amino acid starvation

MDA-MB-231, and SW480 cell lines were used for both hypoxia and amino acid starvation assays.

#### Hypoxia treatment

Cells were seeded in parallel plates and cultured under either normoxic conditions (21% *O*_2_, standard incubator) or hypoxic conditions (1% *O*_2_, 5% *CO*_2_, balance *N*_2_) for approximately 70 h.

#### Amino acid starvation treatment

Amino acid-deficient medium was prepared using DMEM without amino acids (United States Biological) supplemented with dialyzed fetal bovine serum (dFBS). MEM Essential Amino Acids (50× ; Sigma, M5550) and MEM Non-Essential Amino Acids (100× ; Gibco, 11140050) were added to achieve final amino acid concentrations of 40%, or 100% relative to standard medium. MDA-MB-231, and SW480 cells were cultured under these conditions for one day (40% and 100% amino acid media).

Two biological replicates were prepared for each condition and cell line. Total RNA was extracted immediately (Zymo Research) after treatment and used for the construction of the tRNA-seq library using the mature tRNA library mediated by the Y-adapter as previously described (Chen et al., 2025; Shigematsu et al., 2017). The libraries were sequenced as PE80 runs on Nextseq2000 sequencer at Arc Institute.

Raw sequencing reads were processed using `umi_tools` (v1.1.5) to extract unique molecular identifiers (UMIs). Adapter sequences were removed with `cutadapt` (v4.9; parameters: `-e 0.12 -m 20`), and duplicate reads were collapsed based on identical UMI sequences. Paired-end reads were subsequently merged into contiguous fragments with `PEAR` (v0.9.6) and aligned to the human tRNA reference using `bowtie2` (v2.5.4) with the `–very-sensitive` setting. Fragment counts were summarized per tRNA gene, aggregated at the isoacceptor level, and normalized using the `varianceStabilizingTransformation` function implemented in `DESeq2`.

### 5.21. Codon usage analysis of proliferation-, differentiation-, and stress-associated genes

Gene Ontology (GO) terms representing proliferation, differentiation, and tumor stress processes were obtained from the MSigDB GO and Hallmark collections (c5.go.v2024.1.Hs.symbols.gmt) (Liberzon et al., 2015). Proliferation-associated genes were defined by the terms GOBP_CELL_DIVISION and GOBP_MITOTIC_CELL_CYCLE; differentiation-associated genes by GOBP_TISSUE_MORPHOGENESIS and GOBP_PATTERN_SPECIFICATION_PROCESS; and tumor stress-associated genes by GOBP_CELLULAR_RESPONSE_TO_OXIDATIVE_STRESS and GOBP_RESPONSE_TO_NUTRIENT_Genes appearing in different categories were excluded to ensure set-specific comparisons.

For each gene, codon usage frequency was calculated from coding sequences and normalized to the mean frequency of its amino acid encoded codons to obtain synonymous codon usage values. Mean synonymous codon usage frequencies were then calculated across genes within each GO term.

### 5.22. Translational Output Estimation

Matched transcriptome and proteome data from harmonized CPTAC cohorts were used to compute per-gene translational output for each case using a multimodal variational framework inspired by MultiVI (Ashuach et al., 2023). This approach mitigates modality-specific dynamic range differences and systematic measurement effects between RNA and protein, allowing a less biased estimation of post-transcriptional variation.

Specifically, translational output was defined as the component of protein abundance not explained by the shared RNA–protein latent structure. To account for shared biological variation between transcriptome and proteome, we employed a multimodal variational autoencoder that learns (i) a shared latent representation *z*_*s*_ capturing coordinated RNA–protein structure and (ii) a protein-specific latent variable *z*_*p,p*_.

Protein abundance was reconstructed in two modes:

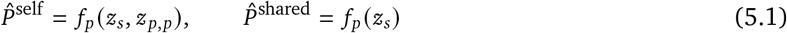

Translational output (TE) was defined as the protein-specific residual:

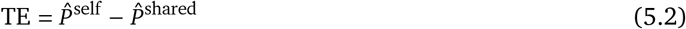

Feature-level TE statistics were computed across samples with observed protein measurements.

### 5.23. Module definition and score calculation

Stress response programs were defined using curated Hallmark and Reactome gene sets. Hallmark and Reactome terms were grouped into biologically related modules, including Hypoxia, Starvation, UPR/ISR, Oxidative ROS, mTOR signaling, Inflammatory NF-κB, and IFN Response. For each module, genes from the corresponding Hallmark or Reactome terms were pooled and deduplicated. In addition, other stress-associated modules were defined, including Mitochondrial Translation, Oxidative Phosphorylation, ER Proteostasis, and ECM Production, as well as proliferation-associated modules, including G2M Checkpoint, E2F Targets, Cell Cycle, and DNA Replication.

Translational output matrices were matched to sample metadata using shared sample identifiers. For each cancer type, translational output values were standardized within cancer by z-scoring each gene across samples of the same cancer type. Genes with zero or near-zero variance within a cancer type were assigned a value of 0 after standardization. Module scores were then computed as the arithmetic mean of available z-scored gene values within each module. Stress-response module scores were used for stress program definition, whereas other stress-associated and proliferation module scores were computed for downstream program activation analysis.

### 5.24. Shared stress extraction

A disentangled variational autoencoder (StressVAE) with separate encoders for shared biological state and cancer identity was used to extract a pan-cancer stress representation that is invariant to cancer type. Module scores were z-scored within cancer prior to training, and all samples were included.

The StressVAE comprised two independent encoders: one generating a shared latent representation and the other generating a cancer-specific latent representation. Each encoder consisted of two fully connected layers with ReLU activation and batch normalization, mapping the input to the mean and log-variance of a low-dimensional Gaussian. These correspond to a shared space (*z*_shared_, dim = 3) capturing pan-cancer stress state, and a cancer-specific space (*z*_cancer_, dim = 4). A joint decoder reconstructed module scores from the concatenation of *z*_shared_ and *z*_cancer_, without direct cancer-type conditioning, forcing *z*_cancer_ to capture identity and cancer-specific environment information.

Training minimized a composite objective:

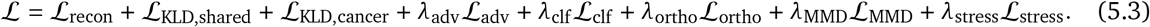

Cancer-type invariance of *z*_shared_ was enforced via a gradient reversal adversary (*λ*_adv_ = 10) and Maximum Mean Discrepancy (MMD) between per-cancer distributions in *z*_shared_ (*λ*_MMD_ = 5, RBF kernel with median bandwidth). A cancer classifier acting on *z*_cancer_ (*λ*_clf_ = 10) encouraged identity concentration in this space. Orthogonality between the two latent spaces was penalized through the mean squared cross-covariance of centered *z*_shared_ and *z*_cancer_ (*λ*_ortho_ = 5).

A stress head—a linear projection from *z*_shared_ to stress module scores—was trained on tumor samples only, beginning at epoch 100 (*λ*_stress_ = 1), providing biological supervision. The adversary was linearly warmed up from epochs 30–100. Models were trained for 300 epochs (Adam, learning rate 1 × 10^−3^, batch size 64) with gradient clipping (maximum norm 5).

### 5.25. Definition of the stress program index

After extraction of the shared stress representation, the mean of the shared latent posterior distribution was obtained for all samples and used as the shared latent embedding. Principal component analysis (PCA) was applied to this representation across all samples, and the first principal component was defined as the stress program index (SPI).

SPI orientation was determined using normal samples as a biological anchor. Specifically, the sign of the first principal component was chosen such that normal samples occupied the lower end of the axis relative to tumor samples. SPI values were then standardized within each cancer type using tumor samples only, generating SPI_*Z*_. For each cancer type, the mean and standard deviation of tumor SPI values were used for scaling, and normal samples from the same cancer type were transformed using the corresponding tumor-derived parameters. For cancer types with sufficient normal samples, tumor displacement from the matched normal centroid was additionally computed in SPI_*Z*_ space.

### 5.26. Parametric stress activation modeling

Parametric activation models were fit separately for each program within each cancer type using SPI_*Z*_ as the independent variable, representing the trajectory of how each molecular program responds along the stress axis. Tumor samples were stratified into up to 20 uniform SPI_*Z*_ bins (minimum 3 samples per bin), and bin-level statistics (median center, mean score, SEM) were used as fitting targets. Inverse-variance weighting was applied throughout.

Three candidate models were considered (Aivazidis et al., 2025):

- **Linear (constant rate):** *f* (*s*) = *c*, representing a program with a constant, stress-independent response rate. A weighted linear trend was additionally computed for visualization and *R*^2^ reporting
- **Sigmoid (threshold):** *f* (*s*) = *b* +*a* ·*σ* (*λ* (*s* −*T*)), representing programs that engage or disengage at a threshold *T* along the SPI_*Z*_ axis.
- **Pulse (transient):** *f* (*s*) = *b*+*a* ·*σ*(*λ*_1_ (*s* −*T*_1_)) ·*σ*(− *λ*_2_ (*s* −*T*_2_)), representing programs transiently activated between SPI_*Z*_ values *T*_1_ and *T*_2_.

Model selection was performed using the Bayesian Information Criterion (BIC). To prevent overfitting of flexible models to approximately linear data, sigmoid and pulse fits were subject to strict validity criteria. Sigmoid fits required: (i) a spanning range of the sigmoidal term ≥ 0.6 across the observed SPI_*Z*_ range; (ii) a threshold *T* within the central 10th–90th percentile of the data (with a ±20% margin); and (iii) a rate concentration ratio ≥ 2.0, defined as the peak absolute response rate divided by its mean.

Pulse fits additionally required: (i) a pulse width between 15% and 80% of the data range; (ii) both *T*_1_ and *T*_2_ within the observed SPI_*Z*_ range; and (iii) a peak amplitude at least twice the residual edge values. Fits failing these criteria were rejected, and BIC selection defaulted to the constant model.

The response rate, defined as the instantaneous derivative 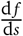 was computed analytically for each model and used to characterize both the stress engagement point and its magnitude.

## 6. Data availability

All sequencing data have been deposited in the Gene Expression Omnibus (GEO) under accession number GSEXXX and will be made publicly available upon publication.

## 7. Acknowledgements

We thank Brian Plosky and Chiara Ricci-Tam for their comments during manuscript preparation. We acknowledge the UCSF Center for Advanced Technology (CAT) and Arc Institute Multi-omics and Labops teams for high throughput sequencing and other experiment supports. We thank the Preclinical Therapeutics core as well as the Laboratory Animal Resource Center (LARC) at UCSF. H.G. is an Arc Core Investigator and research in this paper was partly funded by the Arc Institute. H.G. is an Era of Hope Scholar (W81XWH-2210121) and supported by grants from NCI (R01CA240984 and R01CA244634). V.R. acknowledges funding from NIH grant DP2-HG012442, the Searle Scholars Program, and the W.M. Keck Foundation.

## 8. Author contributions

S.C., H.G. conceived the project. S.C. led the screen and constructed all CRISPRi cell lines; S.C., S.L., T.J. examined the knockdown efficiency of CRISPRi cell lines; S.C. designed tRNA guide sequences; V.R. designed aaRS, tRNA-modifying enzyme, and non-targeting guide sequences; S.C. and T.C. performed the in vitro screen; T.C., K.Y., and B.Z. performed mouse injections and tumor dissections; S.C. optimized the UMI-linked guide RNA library construction protocol; T.C. constructed guide RNA libraries; S.C. performed individual knockdowns, qPCR validation, validation of individual tRNA dependencies under hypoxic conditions, and constructed tRNA libraries under stress conditions. S.C. performed data analyses. S.C. wrote the manuscript; S.C., A.P., V.R., and H.G. revised the manuscript.

